# Organelle Capture, Lineage-Specific Genomic Responses, and the Lability of Dioecy in *Amaranthus*

**DOI:** 10.64898/2026.06.13.732039

**Authors:** David Timerman, Jason Leung, Deren A. R. Eaton

**Affiliations:** Department of Ecology, Evolution and Environmental Biology, Columbia University, New York, NY 10027, USA

**Keywords:** *Amaranthus*, dioecy, cytonuclear evolution, organelle capture, introgression, sex chromosomes, phylogenomics

## Abstract

The transition from combined to separate sexes in plants drives genomic change, from recombination suppression at sex-determining loci to shifts in selection across autosomes and cytoplasmic genomes. In *Amaranthus*, a genus that includes ancient grain crops and noxious weeds, separate sexes arose twice through non-homologous sex-determining architectures, with at least one reversion to monoecy, yet plastid and mitochondrial phylogenies group the two dioecious lineages together despite independent origins. The source of this discordance and whether independent origins of dioecy produced parallel or lineage-specific genomic responses remain unknown. We sampled nuclear, plastid, and mitochondrial genomes across 19 species, resolving the backbone phylogeny of the genus and dating the crown to 2–5 Ma. Coalescent simulations rejected incomplete lineage sorting in favor of multiple organelle capture events, implying historical exchange across reproductive barriers separating the two dioecious clades. Nuclear allele sharing was concentrated within each dioecious clade rather than between them, consistent with recombination eroding nuclear donor ancestry while captured cytoplasmic genomes persist. Dioecy was associated with genome-wide shifts in selection intensity, with positive selection concentrated on the stems of each dioecious clade but targeting largely non-overlapping genes and leaving little signature within sex-determining regions. Plastid coding sequences evolved under relaxed purifying selection, and nuclear-encoded plastid-targeted genes were enriched for episodic positive selection, consistent with compensatory cytonuclear evolution. The probable reversion to monoecy in *A. pumilus*, within a lineage shaped by repeated organelle capture, raises the possibility that hybridization and the lability of separate sexes are connected in this group.

---

*Amaranthus* (Amaranthaceae) is a cosmopolitan angiosperm genus of approximately 70 annual herbs that includes pseudocereal crops, ornamental species, and some of the most problematic agricultural weeds worldwide (Sauer 1955; Waselkov et al. 2018). Species boundaries within the genus have long been difficult to resolve because many taxa are morphologically similar, frequently sympatric, and capable of interspecific hybridization (Sauer 1955; Tranel et al. 2002; Trucco et al. 2005a,b,c). Contemporary gene flow is well documented, including the transfer of herbicide-resistance alleles among weedy species under field conditions and admixture between domesticated grain amaranths and their wild relatives (Gaines et al. 2012; Gonçalves-Dias et al. 2023; Trucco et al. 2005a,b,c, 2009). Despite the ecological and agricultural importance of the genus, phylogenetic relationships across *Amaranthus* remain incompletely resolved, with previous studies based on limited loci or sparse taxon sampling producing conflicting topologies across genomic compartments (Raiyemo and Tranel 2023; Stetter and Schmid 2017; Viljoen et al. 2018; Waselkov et al. 2018; Xu et al. 2024).

Dioecy has evolved at least twice within North American *Amaranthus* lineages, producing approximately nine dioecious species in an otherwise predominantly monoecious genus (Bobadilla et al. 2023; Kreiner et al. 2025; Raiyemo et al. 2025a,b; Waselkov et al. 2018). These species were historically grouped as subgenus *Acnida*, implying a single origin of separate sexes (Mosyakin and Robertson 1996; Sauer 1955). More recent nuclear phylogenies instead place dioecious species into two distinct clades separated by monoecious lineages (Waselkov et al. 2018), and sex-determining regions characterized in representatives of each clade support independent origins through distinct chromosomal locations and nonhomologous genomic architectures (Bobadilla et al. 2023; Kreiner et al. 2025; Montgomery et al. 2021; Neves et al. 2020; Raiyemo et al. 2025a,b). Plastid phylogenies, however, repeatedly place members of these two dioecious lineages closer together than their nuclear positions predict, creating a cytonuclear conflict whose source remains unresolved (Han et al. 2025; Raiyemo and Tranel 2023; Waselkov et al. 2018; Xu et al. 2022).

Conflicts between nuclear and organellar phylogenies can reflect incomplete lineage sorting, introgressive hybridization accompanied by organelle capture, or phylogenetic estimation error (Degnan and Rosenberg 2009; Larson et al. 2026; Rieseberg and Soltis 1991; Wendel and Doyle 1998). Incomplete lineage sorting arises when ancestral polymorphism persists across speciation events, whereas organelle capture records cytoplasmic exchange between diverging lineages (Larson et al. 2026; Tsitrone et al. 2003). Distinguishing among these alternatives requires a multilocus nuclear species tree, coalescent expectations for organellar discordance under incomplete lineage sorting, and independent tests for nuclear introgression (Larson et al. 2026). Because captured cytoplasms are inherited as linked, nonrecombining units while accompanying nuclear ancestry is broken down by recombination, organelle capture can leave a cytoplasmic record of exchange after the nuclear signal has faded (Rieseberg and Soltis 1991; Stull et al. 2023). If the discordance observed in *Amaranthus* reflects this process, the organellar genomes of the genus may preserve a more complete record of historical exchange than the nuclear genome retains.

The independent transitions to dioecy in *Amaranthus* also provide an opportunity to ask whether repeated separation of male and female function produces parallel molecular responses. Dioecy often involves genetic sex determination, and reduced recombination near sex-determining loci can produce differentiated sex chromosomes with divergent gene content (Charlesworth and Charlesworth 1978; Charlesworth 2018; Furman et al. 2020; Leite Montalvão et al. 2021; Marais et al. 2025; Pannell 2017). However, the evolutionary consequences of dioecy can extend beyond sex-determining regions. In cosexual individuals, selection reflects the combined effects of traits on male and female reproductive success within the same organism (Charnov 1982; Lloyd 1982).

Separating male and female function into different individuals can expose alleles to sex-specific selective environments, reveal sexually antagonistic variation to sex-specific selection, allow traits to evolve toward male- and female-specific fitness optima, and alter the efficacy of selection through obligate outcrossing and associated changes in effective population size (Barrett and Hough 2013; Charlesworth 2018; Delph and Ashman 2006; Lande 1980; Lesaffre et al. 2025).

The evolution of dioecy may also alter the selective environment of cytoplasmic genomes. Plastids and mitochondria are predominantly inherited through ovules in flowering plants, so separating male and female function restricts organellar transmission to female lineages (Mogensen 1996). This asymmetry can reduce the effective population size of cytoplasmic genomes and weaken purifying selection, while also allowing male-deleterious organellar variants to persist because they are inefficiently exposed to selection through males, a dynamic known as mother’s curse (Birky et al. 1983; Frank and Hurst 1996; Gemmell et al. 2004; Havird et al. 2019; Postel and Touzet 2020).

Because plastids and mitochondria encode components of molecular complexes whose interacting partners are often nuclear encoded, organellar change may select for compensatory changes in nuclear genes (Postel and Touzet 2020; Sloan 2015; Weng et al. 2016). Hybridization can further perturb this relationship by introducing foreign cytoplasmic haplotypes into new nuclear backgrounds (Shahbazi et al. 2026; Tsitrone et al. 2003). If organelle capture occurred during the history of *Amaranthus*, dioecious lineages may therefore carry cytoplasmic genomes shaped by a different evolutionary history from much of their nuclear genome, creating a potential source of selection at the cytonuclear interface in addition to broader molecular consequences of separate sexes.

Here we use nuclear, plastid, and mitochondrial genomic data from 19 *Amaranthus* species and four outgroup taxa to address four questions. We first ask what the species relationships and divergence times are across the genus, establishing the comparative framework for all subsequent analyses. We then ask whether discordance between nuclear and organellar phylogenies can be explained by incomplete lineage sorting under the nuclear species tree or is better explained by organelle capture through hybridization, testing this explicitly against a coalescent null parameterized from genome-wide nuclear data. We next ask whether the hybridization events inferred from organellar discordance also left detectable signatures in the nuclear genome, or whether recombination has largely erased the nuclear signal while captured organelles persist. Finally, we ask whether the transition to dioecy is associated with shifts in coding-sequence evolution at the cytonuclear interface and in genes relevant to reproductive biology, and whether independent origins of dioecy produce parallel patterns of molecular evolution.

## Materials and Methods

### Taxon Sampling and Sequencing

We assembled a phylogenomic dataset from six annotated *Amaranthus* reference genomes, 31 *Amaranthus* whole-genome short-read accessions representing 19 species, and outgroup resources from public transcriptome, plastome, and short-read datasets. Reference genomes for *A. hybridus*, *A. hypochondriacus*, *A. palmeri*, *A. retroflexus*, *A. tricolor*, and *A. tuberculatus* were obtained from CoGe and NCBI and were selected to span the major clades recovered by Waselkov et al. (2018). Fourteen short-read accessions were newly generated for this study, including 12 USDA GRIN accessions and two field-collected *A. pumilus* accessions; the remaining 17 were drawn from the NCBI Short Read Archive. Outgroups included *Alternanthera philoxeroides*, *Deeringia amaranthoides*, *Hermbstaedtia glauca*, and *Celosia argentea*. Accession identities, provenance, sequencing metadata, and closest-reference assignments are provided in supplementary table S1, Supplementary Material online.

### Phylogenomic Datasets and Trees

#### Nuclear orthogroups

Because outgroup taxa lacked reference genomes, we used a protein-guided framework in which reference proteomes defined the locus set and non-reference samples were added through reference-guided consensus reconstruction. We inferred orthogroups with OrthoFinder v3.1.2 (Deorowicz et al. 2016; Emms and Kelly 2019) from the six *Amaranthus* reference proteomes and three outgroup transcriptome proteomes, retaining orthogroups in which every *Amaranthus* reference contributed exactly one sequence and at least one outgroup sequence was present. We included all outgroup transcript isoforms during clustering rather than collapsing them beforehand, because selecting a representative isoform before orthology inference would require choosing without knowledge of which isoform best matched the conserved coding region of each locus. After clustering, we used twig v0.0.5 (https://github.com/eaton-lab/twig) to remove short and weakly homologous sequences and collapse Trinity isoforms to one representative per outgroup gene. We excluded orthogroups containing sequences with lengths not divisible by three, internal stop codons, or highly uneven lengths across taxa, leaving 4,329 orthogroups.

For each retained orthogroup, we assembled a nucleotide CDS dataset from reference genome annotations, outgroup transcriptome CDS files, and reconstructed sequences from the 31 non-reference *Amaranthus* accessions. We assigned each non-reference accession to its phylogenetically nearest reference genome (supplementary table S1, Supplementary Material online; guided by Waselkov et al. 2018), because coordinate-based CDS reconstruction assumes that exon boundaries are conserved between accession and reference, an assumption more likely to hold for closer relatives. We aligned each accession’s whole-genome reads to its assigned reference with BWA-MEM2 v2.2.1 (Vasimuddin et al. 2019), identified the corresponding gene model in the reference annotation for each orthogroup, extracted per-exon consensus sequences from the accession’s BAM with samtools consensus (minimum depth 5, mapping quality 20, base quality 15), and concatenated those sequences in transcript orientation to yield one reconstructed CDS per accession and orthogroup.

We aligned each orthogroup with twig, which implements a three-stage MACSE v2.07 (Ranwez et al. 2018) codon-aware workflow comprising pre-alignment homology filtering, codon-aware multiple-sequence alignment with reconstructed accession sequences incorporated as lower-confidence references, and post-alignment trimming and overlap filtering. Final alignments were trimmed with ClipKIT v1.3.0 (Steenwyk et al. 2020) using the kpic-smart-gap strategy with codon-aware trimming.

#### Chloroplast genomes

Plastomes for the 31 *Amaranthus* ingroup accessions were assembled with GetOrganelle v1.7.7.1 (Jin et al. 2020), and chloroplast outgroups were downloaded from NCBI (supplementary table S7, Supplementary Material online). Assemblies were processed with PlastidHub v1.0 (Zhang et al. 2025b), standardized to a canonical quadripartite orientation, annotated with PGA v2 (Qu et al. 2019), and used to extract protein-coding genes and intergenic spacers. We excluded protein-coding loci with internal stop codons or other annotation problems and generated a deduplicated dataset retaining one plastome per species for phylogenetic analyses. Coding loci were aligned with the same twig/MACSE codon-aware workflow used for nuclear loci, whereas intergenic spacers were aligned with MAFFT v7.526 (Katoh and Standley 2013). After filtering loci with low taxon occupancy or excessive missing data, we concatenated the remaining loci into a partitioned plastome supermatrix comprising 202 loci (78 protein-coding genes and 124 intergenic spacers).

#### Mitochondrial Loci

Because plant mitochondrial genomes are structurally complex and difficult to assemble from short reads, we recovered individual mitochondrial loci rather than complete mitogenomes. We built a multi-species mitochondrial reference set from published mitochondrial genomes of *A. hypochondriacus, A. palmeri, A. retroflexus, A. tricolor,* and *A. tuberculatus* (supplementary table S8, Supplementary Material online). For each accession, we mapped raw whole-genome reads to a combined reference containing these mitochondrial genomes, the accession’s chloroplast assembly, and its closest nuclear reference genome, then retained read pairs whose strongest mapping support was mitochondrial rather than plastid or nuclear. This competitive-mapping step generated a mitochondria-enriched read set for each sample.

We then mapped the mitochondria-enriched reads for each sample and locus to a locus-specific reference panel containing homologous full-length copies from the curated

*Amaranthus* mitochondrial genomes. For each locus, we selected the panel sequence with dominant read support as the reference for consensus calling and reconstructed consensus sequences with samtools consensus. We excluded loci with insufficient coverage, fragmented or multicopy recovery, excessive ambiguous bases, or internal stop codons. Coding loci were aligned with the same twig/MACSE codon-aware workflow used for nuclear and chloroplast coding loci, whereas non-coding or genomic representations were aligned with MAFFT and trimmed with ClipKIT. After filtering loci with low sample representation, we retained 28 protein-coding loci for phylogenetic analyses (supplementary table S9, Supplementary Material online).

#### Phylogenetic inference

We inferred maximum-likelihood gene trees and organellar trees with IQ-TREE v3.0.1 (Wong et al. 2026), using 1,000 ultrafast bootstrap replicates and BNNI optimization. Nuclear gene trees were inferred under GTR+G and summarized with ASTRAL-IV v1.23.4.6 (Zhang et al. 2025a) to estimate the nuclear species tree under the multispecies coalescent, collapsing multiple accessions per species with an imap file. Trees were rooted on *Alternanthera philoxeroides*, with *Deeringia amaranthoides* and *Hermbstaedtia glauca* used as fallback outgroups when required.

We converted the rooted ASTRAL species tree, pruned to *Amaranthus*, into a chronogram using the penalized-likelihood smoother in toytree v3.1.0.dev1 (Eaton 2020). Alternative clock models were compared with PHIIC and the best-supported model was retained (supplementary table S10, Supplementary Material online). The root age was calibrated from the posterior BPP A00 root *τ* at the deepest sampled *Amaranthus* split (see Coalescent simulation) and converted to absolute time as *T* = *τ*/*μ*. Because no direct *Amaranthus* coding-sequence mutation rate is available, we report ages under the 7 × 10^−9^ substitutions site^−1^ year^−1^ Arabidopsis whole-genome reference clock (Ossowski et al. 2010), assuming a one-year generation time, and under a more conservative rate of 2.94 × 10^−9^ substitutions site^−1^ year^−1^ derived by scaling the reference by the 58% lower mutation frequency observed within coding sequences of *Arabidopsis* (Monroe et al. 2022). We then reconstructed ancestral sexual-system states on the chronogram with a two-state equal-rates continuous-time Markov model in toytree.pcm, coding species as monoecious or dioecious and fixing the root prior to monoecious. We assessed uncertainty with 200 stochastic-character maps under the fitted model.

For chloroplast data, we inferred maximum-likelihood trees from full and deduplicated plastome supermatrices under ModelFinder model selection. For mitochondrial data, we inferred the primary tree from the partitioned protein-coding matrix and evaluated alternative partitioning and sequence-treatment schemes as sensitivity analyses. Organellar trees were rooted on *A. philoxeroides*.

### Coalescent Simulations

We tested whether incomplete lineage sorting alone could explain organellar discordance by simulating organellar genealogies under the nuclear species tree. Coalescent parameters were estimated with fixed-topology BPP v4.8.0 A00 analyses (Flouri et al. 2018) on filtered nuclear orthogroup alignments (500–1,200 bp, ≥ 300 high-quality sites per sequence), using the rooted *Amaranthus* ASTRAL tree as the fixed topology. BPP estimated divergence times (*τ*) and population mutation rates (*θ*) under GTR, with inverse-gamma priors calibrated from pairwise sequence divergence and mean per-site heterozygosity. We used posterior median *τ* and *θ* estimates to parameterize ipcoal v0.5.0 simulations (McKenzie and Eaton 2020) on the nuclear species tree. To approximate haploid, uniparentally inherited organellar genomes, we rescaled diploid nuclear *θ* estimates by 1/16, converting *θ* = 4*N*_e_*μ* to *N*_e_*μ* and then applying a 0.25 organellar inheritance scaling. We simulated 10^6^ independent, non-recombining genealogies with one lineage per tip.

We compared the observed chloroplast and mitochondrial trees to the simulated null distribution in two ways. First, we compared their normalized information-corrected Robinson-Foulds (RFI) distances from the nuclear species tree to the distribution of simulated genealogies. Second, for each non-trivial bipartition in the observed organelle trees, we calculated the fraction of simulated genealogies containing the same bipartition, yielding a null probability for each observed split. As a complementary per-locus comparison, we computed ΔRFI as the distance of each nuclear gene tree to an organelle tree minus its distance to the species tree.

### Phylogenetic Network and Admixture Analyses

To characterize historical gene flow, we combined nuclear gene-tree network inference, SNP-based admixture tests, and coalescent model comparisons. We inferred reticulate relationships from nuclear gene trees with PhyloNet v3.8.2 (Than et al. 2008; Wen et al. 2018) under the maximum pseudo-likelihood framework (Yu and Nakhleh 2015), running 1,000 independent searches for each reticulation count (k = 0–5) and retaining the five highest-scoring networks. We selected the best network for each k and identified a parsimonious reticulation count from the elbow in log pseudo-likelihood improvement.

For SNP-based tests, we generated a joint biallelic SNP dataset from 30 *Amaranthus* accessions plus *A. philoxeroides* by aligning reads to the *A. tricolor* reference with BWA-MEM2, processing alignments with samtools v1.23 (Danecek et al. 2021), and calling variants with BCFtools v1.20 (Danecek et al. 2021). We retained biallelic SNPs after filtering for minimum mapping quality 20, minimum site quality 25, maximum missing data 20%, minimum distance from indels 5 bp, maximum depth twice the genome-wide mean, and minimum minor allele count 2. We analyzed the resulting dataset with Dsuite v0.5 r58 (Malinsky et al. 2021), assigning accessions to 19 *Amaranthus* species with *A. philoxeroides* as outgroup. We computed Patterson’s D (Durand et al. 2011), the *f*_4_-ratio, and *F*_branch_ across all 969 species trios, assessing significance from block-jackknife *Z*-scores under the conservative Dmin arrangement with Benjamini-Hochberg false-discovery-rate correction and using tree-based trio orientations for directional D-statistics and *f*_4_-ratios. We tested for clustered ABBA-informative sites with Dsuite –ABBAclustering (Koppetsch et al. 2024) and localized signals with Dinvestigate analyses in 50-SNP windows with 25-SNP steps.

Finally, we estimated the direction and magnitude of candidate introgression events with quartet-level BPP v4.8.7 MSci analyses (Flouri et al. 2020). For each quartet, we retained loci containing all four target species, at least 250 unambiguous sites per sequence, and alignment lengths of 200–1,000 bp. We compared four models (no introgression, introgression in either direction, and bidirectional introgression) using the same substitution model and *τ* and *θ* priors as the A00 analysis, a Beta(1,1) prior on introgression proportion *ϕ*, and two independent MCMC chains per model with 20,000 burn-in generations and 100,000 post-burn-in samples thinned every five generations.

### Selection Analyses

We applied three HyPhy v2.5.96 methods to test for coding-sequence shifts associated with dioecious lineages across 4,329 deduplicated nuclear codon alignments and their gene trees, applying RELAX additionally to concatenated chloroplast and mitochondrial protein-coding alignments. RELAX (Wertheim et al. 2015) estimates a selection-intensity parameter K by comparing distributions of *ω* (*d*_*N*_ /*d*_*S*_) across site classes on foreground and reference branches. BUSTED-PH (Selberg et al. 2026) tests whether foreground branches exhibit an elevated proportion of sites evolving under episodic positive selection (*ω* > 1) relative to reference branches. aBSREL (Smith et al. 2015) tests individual branches for episodic diversifying selection without requiring a predefined foreground-reference partition.

For RELAX and BUSTED-PH, we assigned branch labels by sexual system using a conservative conjunctive rule: terminal branches were labeled by taxon sexual system and internal branches were labeled only when all descendant taxa belonged to the same class, with mixed or ambiguous branches left unspecified. We defined two nuclear contrasts, Dioecious I+II and Dioecious I alone, leaving outgroups, as well as monoecious *A. pumilus* and *A. spinosus* unspecified in both; we did not define a Dioecious II-only contrast because only two dioecious species were sampled from that clade. The organellar RELAX analyses used the combined Dioecious I+II foreground.

Because BUSTED-PH and aBSREL are sensitive to alignment and annotation errors that can mimic episodic positive selection, we applied the BUSTED-E error-filter pipeline (Selberg et al. 2025) before both analyses. BUSTED-E models an error-sink class for codon patterns more consistent with sequencing or annotation artifacts than with genuine selection, and flagged sites were masked before downstream analyses. RELAX was applied to unfiltered alignments because it estimates shifts in selection intensity rather than episodic positive selection, and the error-sink model is not parameterized for that purpose. For aBSREL, we retained branches at corrected *p* ≤ 0.05 that passed numerical quality filters, excluding branches with fewer than 0.001 synonymous substitutions per site or length below 0.01 and models in which *ω* reached extreme values indicative of numerical instability.

We annotated nuclear loci using the *A. hypochondriacus* v3 annotation of Graf et al. (2025), supplemented by *Arabidopsis thaliana* reciprocal-best DIAMOND v2.1.25 hits against Araport11 representative proteins (Buchfink et al. 2021). Plastid and mitochondrial targeting of nuclear-encoded proteins was assessed from integrated TargetP v2.0 (Almagro Armenteros et al. 2019) and DeepLoc v2.1 (Ødum et al. 2024) predictions. We tested enrichment of organelle-associated proteins among candidate gene sets with one-sided Fisher exact tests and Benjamini-Hochberg FDR correction, tested GO-term enrichment with GOATOOLS v1.6.4 (Klopfenstein et al. 2018), and screened candidates against curated sexual-system GO categories, including reproductive and floral development, fertility, pollination and pollen-tube biology, meiosis and recombination, hormone signaling, and transcriptional and epigenetic regulation (supplementary table S6, Supplementary Material online).

## Results

### Fully Resolved Nuclear Backbone and Sexual-System Transitions

The nuclear dataset comprised 4,329 single-copy coding alignments from 37 accessions representing 19 *Amaranthus* species and three outgroups, with short-read accessions contributing reconstructed coding sequences at median 13.2× coverage and outgroup transcriptomes contributing sequences to 87–91% of loci (supplementary tables S1 and S2, Supplementary Material online).

The species tree was fully resolved, with local posterior probability of 1.0 at nearly every internal branch, and recovered four major clades (Early-diverging, Dioecious I, Dioecious II, and Hybridus/grain; Fig. 1). The Early-diverging clade, comprising *A. fimbriatus*, *A. viridis*, and *A. tricolor*, was sister to the rest of the genus. The remaining species split into Dioecious I and a clade uniting Dioecious II with Hybridus/grain, with full support at both internal branches.

**Figure 1:**
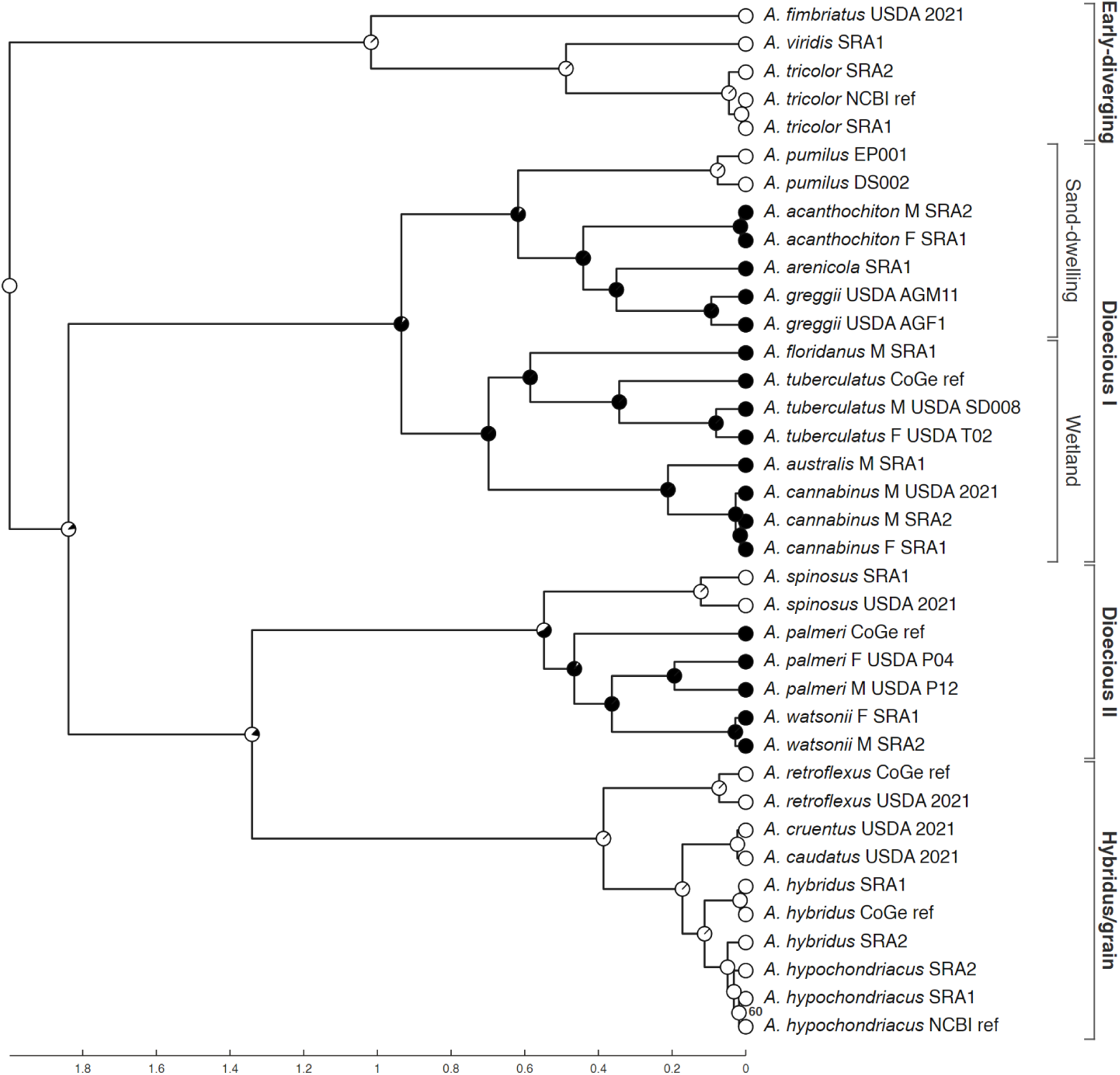
Nuclear species tree of *Amaranthus*. Rooted ASTRAL species tree converted to a chronogram with branch lengths in millions of years, calibrated to the 7 × 10^−9^ substitutions site^−1^ year^−1^ reference clock (reflecting a crown-age range of 2–5 Ma under the two clock rates considered). Tip markers denote sexual system: filled, dioecious; open, monoecious. Internal labels indicate local posterior probabilities below 0.95; unlabelled branches exceed 0.95. Right-side brackets mark major clades and groups used in downstream analyses. Alt text: Time-calibrated phylogenetic tree of 19 *Amaranthus* species with branch lengths in millions of years. Branches are organized into four labeled clades. Filled tip circles indicate dioecious species and open circles indicate monoecious species. Pie charts at internal nodes show posterior probabilities of ancestral sexual-system states. Internal branch labels show local posterior probability values below 0.95.

Dioecious I contained seven dioecious species and the monoecious *A. pumilus*, with *A. pumilus* sister to a sand-dwelling subclade comprising *A. acanthochiton*, *A. arenicola*, and *A. greggii*, and the sand-dwelling subclade in turn sister to a wetland subclade comprising *A. floridanus*, *A. tuberculatus*, *A. australis*, and *A. cannabinus*. Dioecious II contained the dioecious *A. palmeri* and *A. watsonii* together with the monoecious *A. spinosus*, and was sister to Hybridus/grain, which contained *A. retroflexus* and the grain amaranths *A. caudatus*, *A. cruentus*, *A. hybridus*, and *A. hypochondriacus*. *A. watsonii* fell within *A. palmeri* and *A. hypochondriacus* fell within *A. hybridus*, but neither pattern altered relationships among the major clades.

The posterior BPP root *τ* at the deepest sampled *Amaranthus* split was 0.01399 (95% HPD 0.01369–0.01429), corresponding to a crown-age range of 2–5 Ma under the two clock rates applied (Fig. 1). Ancestral state reconstruction recovered two independent origins of dioecy corresponding to the Dioecious I and Dioecious II clades. Within Dioecious I, *A. pumilus* was confidently reconstructed as a reversion to monoecy nested in the otherwise-dioecious sand-dwelling subclade. Within Dioecious II, the ancestral state at the crown node was ambiguous (*P*_dioecious_=0.56, *P*_monoecious_=0.44), leaving unresolved whether dioecy arose once at the Dioecious II crown with subsequent reversion in *A. spinosus* or independently on the *A. palmeri* + *A. watsonii* branch (*P*_dioecious_=0.99), with *A. spinosus* retaining ancestral monoecy.

### Cytonuclear Discordance and Evidence for Organelle Capture

The chloroplast dataset comprised 202 loci (78 protein-coding genes and 124 intergenic spacers; 130,621 bp; 33 tips) and the mitochondrial dataset comprised 28 protein-coding loci (35,120 bp; 21 tips). Both organellar phylogenies were fully resolved, but support was stronger and more uniform in the chloroplast tree, where all but one internal branch had UFBoot > 95, than in the mitochondrial tree (Fig. 2). Both conflicted strongly with the nuclear species tree, and the largest rearrangements were shared across compartments.

**Figure 2:**
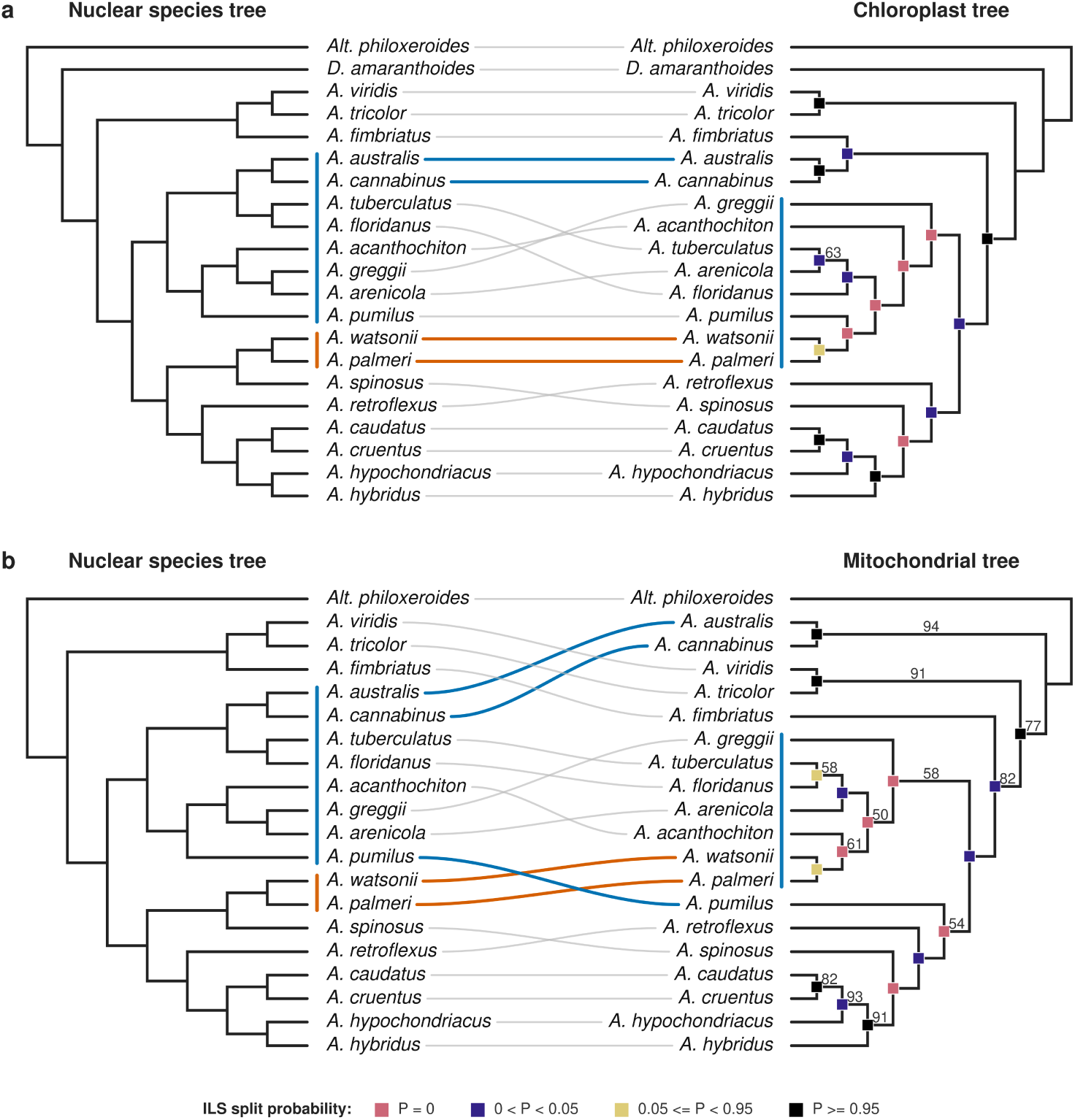
Nuclear-organelle topology discordance in *Amaranthus*. Tanglegrams compare the nuclear ASTRAL species tree with the chloroplast (A) and mitochondrial (B) trees. Lines connect matching taxa; colored bars mark Dioecious I and the *A. palmeri*/*A. watsonii* lineage, with discordant taxa highlighted by colored connections. Squares on organelle branches indicate the probability of recovering each split under an ILS null model scaled to organelle effective population size. Alt text: Two tanglegram panels, each pairing the nuclear species tree on the left with an organellar tree on the right. Panel A shows the chloroplast tree and panel B shows the mitochondrial tree. Colored lines connect taxa whose organellar positions conflict with their nuclear positions, concentrated in the Dioecious I and Dioecious II clades. Squares on organellar branches indicate ILS null probabilities.

In both organellar trees, Dioecious II was polyphyletic. *A. palmeri* and *A. watsonii* were displaced from their nuclear position with *A. spinosus* and instead grouped with members of Dioecious I, while *A. spinosus* grouped with Hybridus/grain. The two compartments agreed on the displacement of *A. palmeri* and *A. watsonii* into Dioecious I but differed in the specific Dioecious I lineage they associated with; in the chloroplast tree they were sister to *A. pumilus* within the sand-dwelling subclade, and in the mitochondrial tree they were sister to *A. acanthochiton*. *A. spinosus* nested within Hybridus/grain in both trees. *A. pumilus* was also displaced from its nuclear position within Dioecious I in the mitochondrial tree, appearing sister to Hybridus/grain, though this alternative placement was not strongly supported.

A second major discordance involved the wetland lineage containing *A. australis* and *A. cannabinus*. In the chloroplast tree these species grouped with *A. fimbriatus* as sister to the rest of Dioecious I, Dioecious II, and Hybridus/grain. In the mitochondrial tree they occupied the earliest-diverging position within *Amaranthus*, with *A. fimbriatus* sister to the remaining ingroup. Both compartments separated the wetland lineage from its nuclear position within Dioecious I.

Posterior BPP estimates of *θ* varied by more than two orders of magnitude across terminal branches, from 4.8 × 10^−5^ in *A. hypochondriacus* to 0.020 in *A. palmeri*, and *τ* at the deepest *Amaranthus* split was 0.01399 (95% HPD 0.01369–0.01429; supplementary table S3, Supplementary Material online). Coalescent simulations scaled to organellar inheritance placed both observed organellar trees far outside the null distribution; the observed chloroplast and mitochondrial RFI distances of 0.789 and 0.756 exceeded every simulated genealogy (*p* < 10^−6^; Fig. 3). The excess discordance was concentrated in a discrete set of splits rather than distributed evenly across the trees. In the chloroplast tree, 11 of 17 non-trivial bipartitions had null probabilities below 0.05 and five were never recovered in 10^6^ simulations, with the most extreme splits linking *A. palmeri* and *A. watsonii* to *A. pumilus*. The mitochondrial tree exhibited a similar pattern, with 10 bipartitions below *p* = 0.05 and five never recovered, with the most extreme splits linking *A. palmeri* and *A. watsonii* to *A. acanthochiton*. The displaced positions of *A. australis* and *A. cannabinus* produced additional zero-probability splits in both compartments. These results reject incomplete lineage sorting as a sufficient explanation for the organellar topologies and support organelle capture as the source of the major cytonuclear discordance.

**Figure 3:**
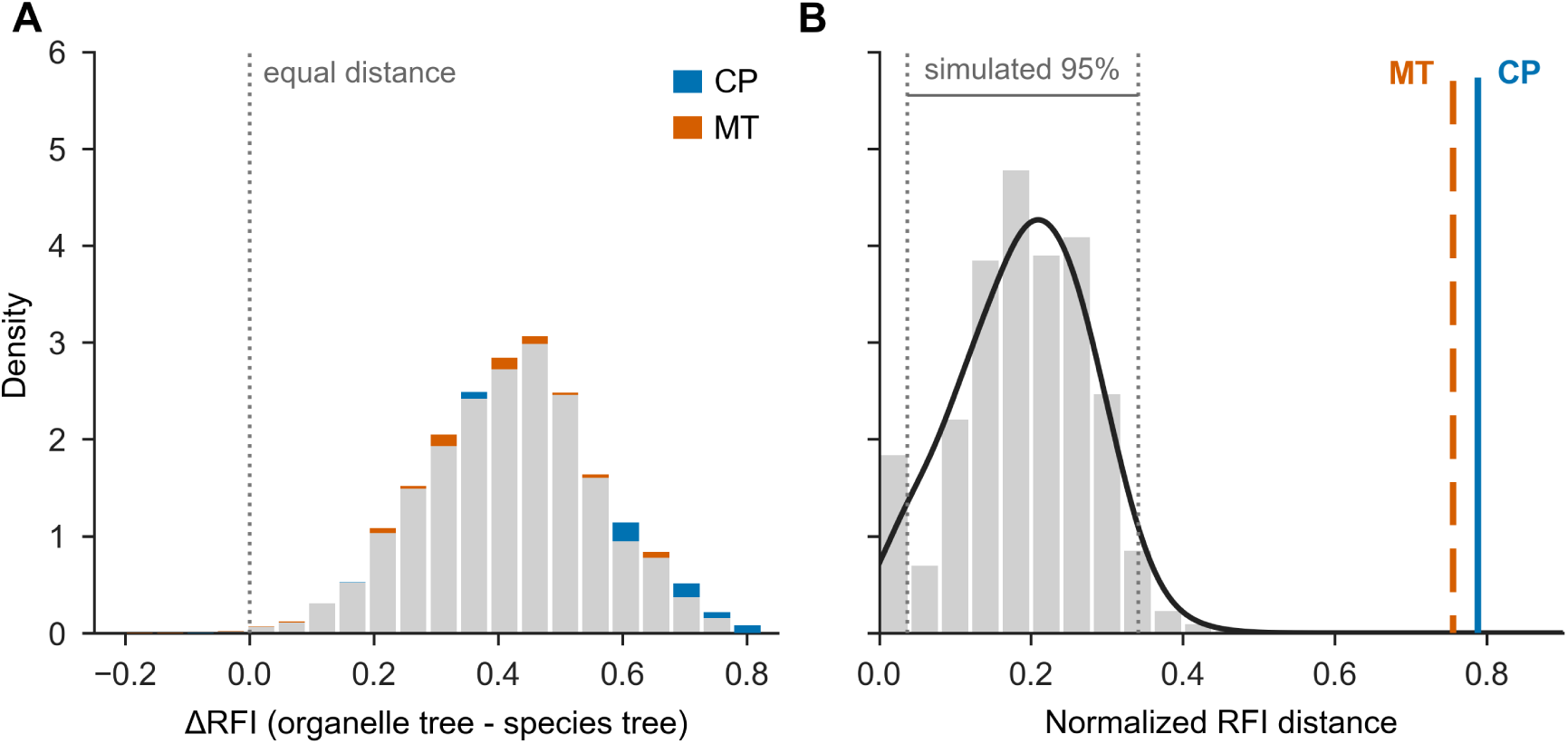
Organelle topology discordance. (A) Per-locus difference between each nuclear gene tree’s normalized Robinson–Foulds information distance to the chloroplast or mitochondrial tree and its distance to the nuclear species tree (positive values indicate greater similarity to the species tree). Gray segments indicate per-bin overlap between organelles; colored caps indicate excess for either organelle. (B) Null distribution of distances from ILS-simulated gene trees to the species tree, with observed chloroplast and mitochondrial distances overlaid. Distances are normalized from 0, identical topology, to 1, maximum discordance. Alt text: Two-panel figure. Panel A is a histogram showing per-locus Robinson-Foulds distance differences between organellar and nuclear topologies, with chloroplast and mitochondrial distributions shown in overlapping colored segments. Panel B shows a histogram of ILS-simulated null distances with observed chloroplast and mitochondrial distances marked as vertical lines falling far in the right tail of the distribution.

### Reticulate Evolution Across Multiple Evolutionary Timescales

PhyloNet maximum pseudo-likelihood analyses identified a three-reticulation network as the most parsimonious summary of the nuclear gene-tree distribution (Fig. 4A). Model fit improved steeply from k = 0 to k = 3, with successive log pseudo-likelihood gains of 42,193, 23,603, and 9,239, compared with 2,740 and 1,210 at k = 4 and k = 5. The first reticulation localized to Dioecious I, where the *A. floridanus*/*A. tuberculatus* lineage was recovered as the reticulation descendant in 65.3% of 1,000 independent k = 1 searches. A second signal involving *A. fimbriatus* appeared at k = 2 and strengthened at k = 3, where its most frequent tripartition, recovered in 24.4% of searches, placed it between the *A. tricolor* + *A. viridis* clade and the Dioecious II + Hybridus/grain lineage. The best k = 3 network resolved two recurrent reticulation regions, one involving *A. fimbriatus* near the early-diverging backbone and one within Dioecious I comprising two nested reticulations centered on *A. floridanus*, *A. tuberculatus*, and neighboring lineages. In the Dioecious I reticulations, the donor was placed consistently among sand-dwelling rather than wetland species, and the mean minor-parent inheritance probability across the three reticulations was approximately 0.26.

**Figure 4:**
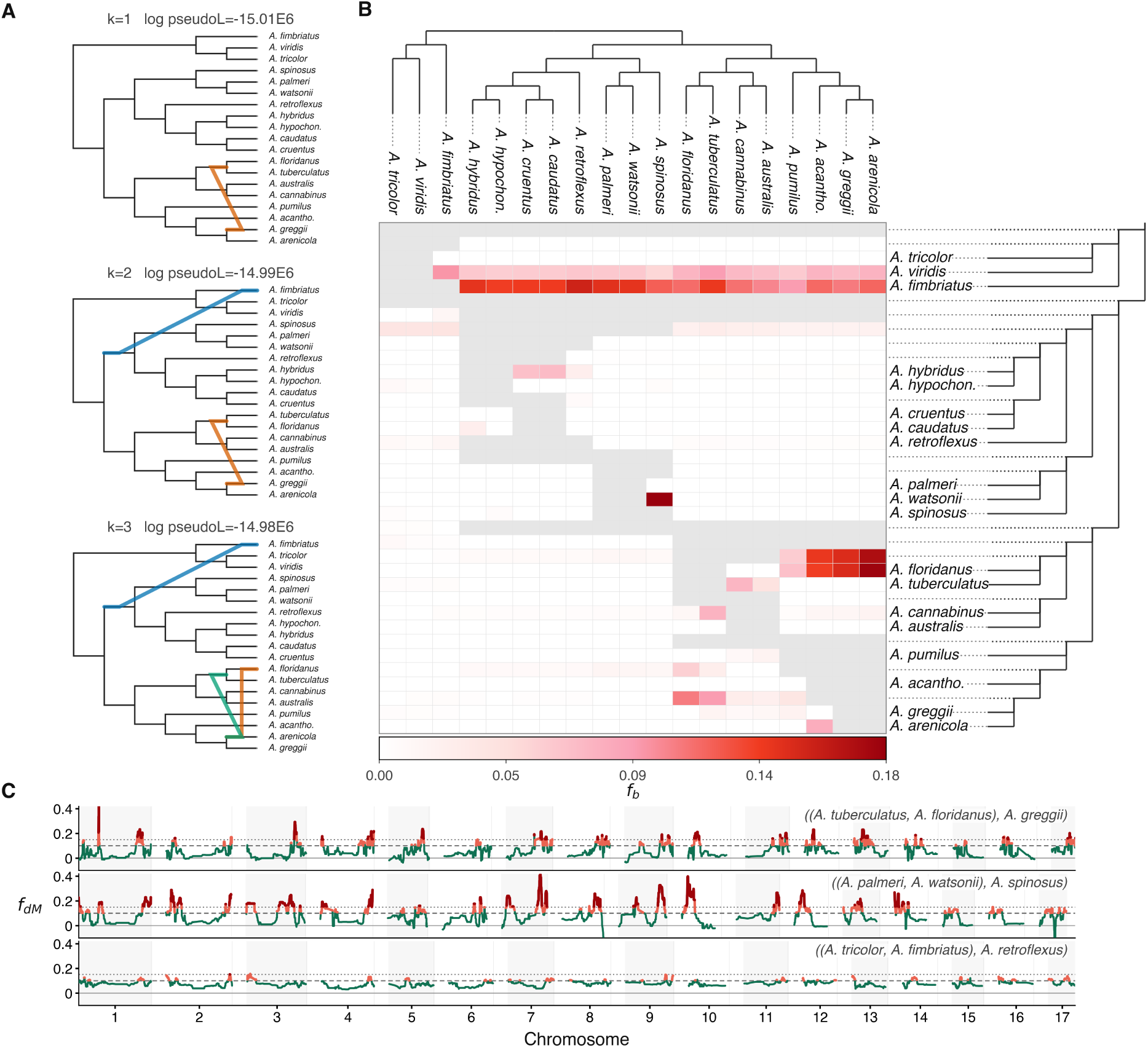
Reticulate evolution in the *Amaranthus* nuclear genome. (A) PhyloNet maximum pseudo-likelihood networks inferred from nuclear gene trees with one, two, and three reticulations; edge colors indicate the network in which each signal first appears. (B) Dsuite *F*_branch_ heatmap of excess allele sharing across the nuclear genome. Gray cells indicate unevaluable comparisons. (C) Windowed *f*_*dM*_ tracks for three tests of the form ((P1, P2), P3) across the *A. tricolor* reference genome. Green lines indicate rolling medians; red overlays mark *f*_*dM*_ ≥ 0.10 and *f*_*dM*_ ≥ 0.15. Outgroup: *Alternanthera philoxeroides*. Alt text: Three-panel figure. Panel A shows phylogenetic networks with one, two, and three reticulation events depicted as colored reticulate edges on a species tree backbone. Panel B shows a species-by-species heatmap matrix of excess allele sharing values, with darker cells indicating stronger signals in four phylogenetic regions. Panel C shows three genome-wide tracks of windowed introgression statistics plotted along the *A. tricolor* reference chromosomes, with rolling median lines and highlighted regions of elevated signal.

Joint variant calling against the *A. tricolor* reference yielded 11,099,793 biallelic SNPs across 30 *Amaranthus* accessions and *Alternanthera philoxeroides* as outgroup (supplementary table S4, Supplementary Material online). Across the 969 trios among the 19 species, Patterson’s D was significant in 758 trios after Benjamini-Hochberg correction, but *F*_branch_ concentrated the strongest signals into four phylogenetic regions (Fig. 4B). Two regions corresponded to the PhyloNet reticulations. The early-diverging component involved elevated allele sharing between *A. fimbriatus* and most non-early-diverging lineages (*F*_branch_ up to 0.154, *f*_4_-ratios up to 0.183); within the early-diverging clade, *A. viridis* exhibited a similarly broad but weaker pattern (*F*_branch_ 0.052–0.095, *f*_4_-ratios up to 0.126 in trios contrasting *A. viridis* and *A. fimbriatus*), while *A. tricolor* did not. The Dioecious I component linked the wetland lineage with sand-dwelling *A. arenicola*, *A. greggii*, and *A. acanthochiton* (*F*_branch_ 0.171–0.179), matching the PhyloNet reticulation zone. The third component was a localized signal between *A. watsonii* and *A. spinosus* (*F*_branch_ = 0.182), and a fourth weaker component appeared within Hybridus/grain, with elevated allele sharing between *A. hybridus* and the *A. cruentus*/*A. caudatus* side of the clade (*F*_branch_ 0.068–0.071).

Windowed Dinvestigate distinguished tract-like introgression from diffuse allele sharing (Fig. 4C). In Dioecious I, tract structure was strongest in contrasts between wetland and sand-dwelling lineages, with *A. floridanus* sharing excess derived alleles with *A. greggii* over *A. tuberculatus* (mean *f*_*dM*_ = 0.086, longest positive run 1.45 Mb; Fig. 4C, top). Reciprocal contrasts of *A. floridanus* and *A. tuberculatus* against *A. arenicola* gave similar *f*_*dM*_ values and passed robust ABBA-clustering tests (*p* ≤ 0.02). Trios involving *A. pumilus* were consistently weaker. The strongest tract structure in any contrast linked *A. watsonii* with *A. spinosus* (*D* = 0.259, *Z* = 16.4, *f*_4_-ratio = 0.182, robust ABBA-clustering *p* = 0.019, mean *f*_*dM*_ = 0.115, longest positive run 2.93 Mb spanning 41 consecutive windows; Fig. 4C, middle). Quartet-level BPP MSci analyses corroborated bidirectional introgression between *A. watsonii* and *A. spinosus*, rejecting the no-admixture tree (Δ ln *L* ≈ 2, 000), with estimated introgression fractions of 0.14 (95% interval 0.130–0.156) for *A. spinosus* → *A. watsonii* and 0.47 (95% interval 0.438–0.502) for *A. watsonii* → *A. spinosus* (supplementary table S5, Supplementary Material online).

In contrast, the early-diverging and Hybridus/grain signals were strong genome-wide but weakly structured across windows. Within the early-diverging clade, *A. fimbriatus* shared excess derived alleles with *A. viridis* over *A. tricolor* (*D* = 0.364), but the local signal was diffuse (mean *f*_*dM*_ = 0.045, 3.25% of windows above *f*_*dM*_ = 0.1). *A. fimbriatus* also shared elevated derived alleles with *A. retroflexus* over *A. tricolor*, but this signal was similarly diffuse (mean *f*_*dM*_ = 0.080, longest positive run 0.89 Mb; Fig. 4C, bottom). None of the 49 trios involving early-diverging lineages passed the robust ABBA-clustering test, and BPP quartet models found no evidence of recent direct exchange between *A. fimbriatus* and either *A. viridis* or *A. retroflexus*. The only chain-concordant directional model estimated an introgression fraction near zero (1 − *ϕ* = 0.001, 95% interval 0.000–0.004). The Hybridus/grain signal exhibited a similar profile, with allele sharing distributed across many short windows rather than concentrated in long tracts.

### Coding-Sequence Evolution on Dioecious Branches

RELAX identified 47 FDR-significant loci with evidence of shifted selection intensity in the Dioecious I contrast, nearly evenly split between relaxation (*n* = 24) and intensification (*n* = 23), and 44 loci in the combined Dioecious I+II contrast (Fig. 5A). BUSTED-PH identified 16 loci with elevated episodic positive selection in Dioecious I and 13 in the combined contrast, all of which were also significant in the Dioecious I-only analysis. Branch-level aBSREL identified 280 loci with branches revealing episodic positive selection after quality filtering, most of which were internal rather than terminal; 185 of 253 Dioecious I branch hits occurred on internal branches subtending the sampled clade, and 40 of 54 Dioecious II-neighborhood hits occurred on the stem branch of the *A. palmeri*, *A. watsonii*, and *A. spinosus* clade (Fig. 5B). Only 10 loci carried aBSREL-supported branches in both dioecious branch sets.

**Figure 5:**
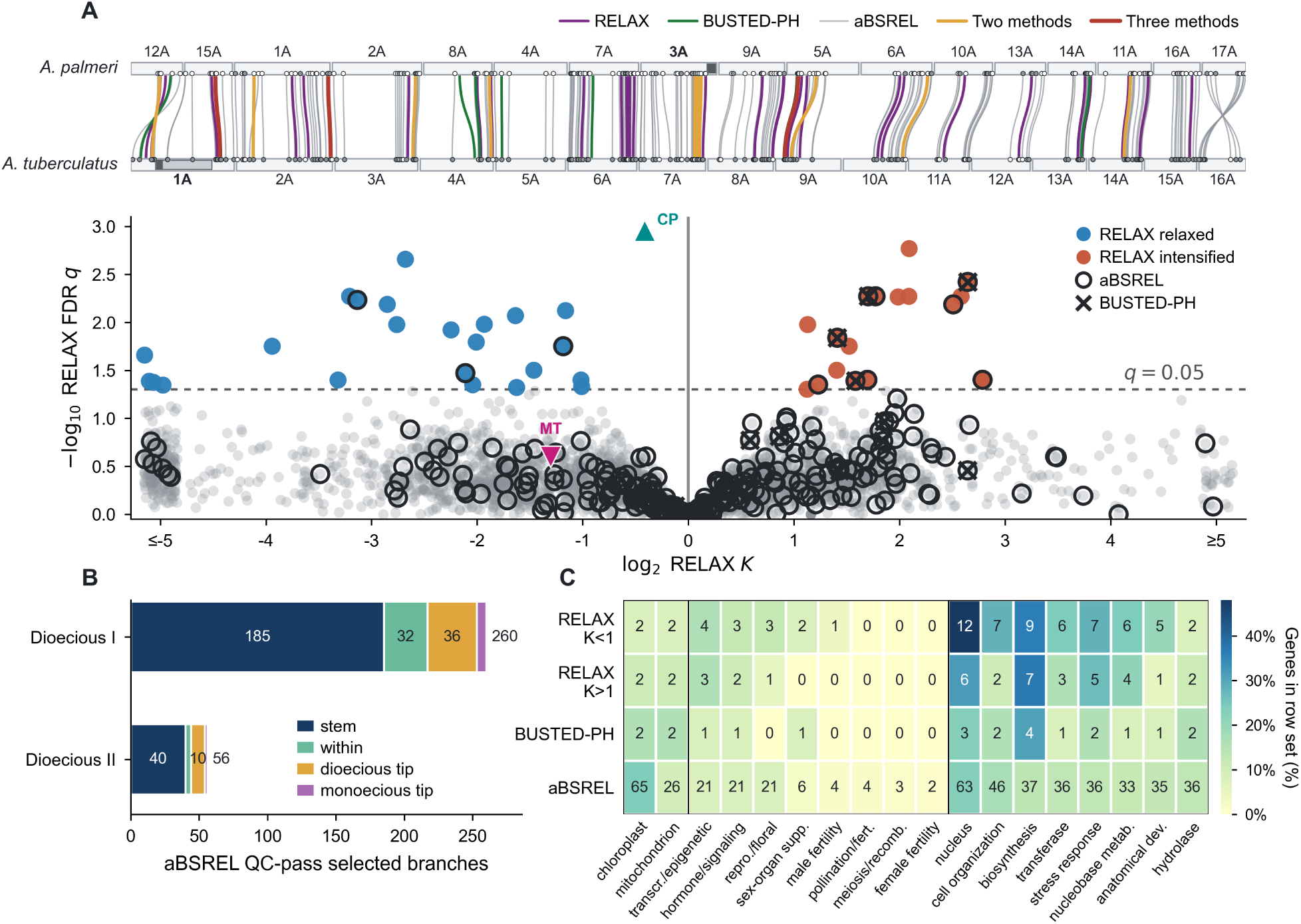
Selection signatures associated with dioecy in *Amaranthus*. (A) Upper, synteny map of single-copy nuclear orthogroups between the *A. palmeri* and *A. tuberculatus* reference genomes. Ribbons are colored by HyPhy method for loci significant in the Dioecious I contrast (RELAX and BUSTED-PH) or in either dioecious clade (aBSREL); putative sex-determining regions are shaded. Filled endpoint circles mark loci with aBSREL-significant selection on the stem leading to that reference’s clade (Dioecious I for *A. tuberculatus*; Dioecious II for *A. palmeri*); open circles mark the non-significant stem on that side. Lower, RELAX results for the Dioecious I+II contrast, with loci also recovered by aBSREL or BUSTED-PH marked by overlaid symbols. Dashed line marks *q* = 0.05; log_2_ *K* is clipped at ±5. (B) QC-pass aBSREL-selected branches in each dioecious lineage, partitioned by branch class; embedded monoecious tips are *A. pumilus* and *A. spinosus*. (C) Functional features across selected gene sets, including organellar targeting, curated dioecy-relevant GO categories, and frequent GO-slim context terms. Alt text: Three-panel figure. Panel A shows a synteny river plot linking orthologous loci between the *A. palmeri* and *A. tuberculatus* reference genomes, with ribbons colored by HyPhy analysis method and putative sex-determining regions shaded, plus a strip below showing RELAX selection-intensity values along the genome. Panel B shows a stacked bar chart of aBSREL-selected branches partitioned by branch class for each dioecious lineage. Panel C shows a dot plot of functional annotations across candidate gene sets, with dot size or shading indicating frequency of GO terms and organellar targeting categories.

Significant loci were distributed across the genome rather than concentrated in the characterized sex-determining regions (Fig. 5A), though some occurred on chromosomes carrying sex-determining regions, including XRCC4, a DNA repair factor, and EMB2771, annotated for cell-cycle regulation and female fertility, both on *A. tuberculatus* chromosome 1; none mapped to the core regions themselves. Most significant loci were identified by only one of the three tests, but 29 in Dioecious I and 21 in the combined contrast were supported by multiple methods. Five loci were significant across all three methods in both contrasts, including TPS-CIN, a terpene synthase, THO1, a subunit of the mRNA export complex, and FLAP1, a nuclear-encoded chloroplast-targeted protein. The monoecious species *A. pumilus* and *A. spinosus* exhibited no excess of terminal-branch aBSREL hits relative to neighboring dioecious taxa, though the seven terminal-branch hits in *A. pumilus* included RMI2, a RecQ-associated genome-stability factor annotated for meiotic crossover dissolution and pollen development.

No functional class except plastid-associated genes passed FDR-corrected enrichment. A curated screen of dioecy-relevant GO categories identified 56 of 280 loci with aBSREL-supported branches associated with reproductive or floral development, pollination and fertilization, meiosis and recombination, hormone signaling, or transcriptional and epigenetic regulation (supplementary table S6, Supplementary Material online), including CLV2, a CLAVATA-pathway receptor involved in floral meristem regulation and reported as differentially expressed between male and female flowers in *A. palmeri* (Bobadilla et al. 2023). Nuclear-encoded plastid-targeted proteins were enriched among loci with aBSREL-supported branches; in the combined Dioecious I+II branch set, 65 of 272 loci were plastid-associated compared with 746 of 4,317 background loci (23.9% versus 17.3%; 1.38-fold enrichment; *q* = 0.032; Fig. 5C). Plastid-associated genes were also overrepresented among stem-branch hits in both clades. RELAX and BUSTED-PH loci showed no parallel plastid enrichment, and mitochondrial-associated loci were not significantly enriched in any test.

Chloroplast protein-coding sequences evolved under relaxed purifying selection on dioecious foreground branches (*K* = 0.750, *LRT* = 10.64, *p* = 0.0011), with the strongest per-branch support concentrated in Dioecious I, including *A. cannabinus* (*E R* = 229), *A. australis* (*E R* = 107), their shared ancestral branch (*E R* = 12.3), *A. acanthochiton* (*E R* = 11.1), and the branch subtending *A. floridanus*, *A. arenicola*, and *A. tuberculatus* (*E R* = 7.66). *A. cannabinus* and *A. australis*, whose chloroplast positions are displaced outside Dioecious I in the organellar trees (Fig. 2), carry captured plastomes and showed among the strongest per-branch relaxation signals in the dataset. Mitochondrial coding sequences showed no significant shift in selection intensity (*K* = 0.405, *LRT* = 1.32, *p* = 0.251).

## Discussion

We present a new genome-scale nuclear phylogeny of *Amaranthus*, resolving the source of cytonuclear discordance in the genus as ancient organelle capture, and identifying plastid-linked shifts in coding-sequence evolution associated with dioecious lineages. The nuclear tree resolved the backbone and rooting of the genus, supported two independent origins of dioecy, and identified at least one likely reversion to monoecy. Comparisons with chloroplast and mitochondrial trees revealed that organellar discordance reflected ancient cytoplasmic exchange rather than incomplete lineage sorting, while nuclear network and allele-sharing analyses recovered a broader history of introgression that did not simply mirror the organelle captures. Selection analyses revealed that dioecious lineages carried shifts in coding-sequence evolution consistent with the expectation that separate sexes should alter selective pressures across the genome, including concentrated episodic selection on dioecious-clade stems but little overlap in the specific genes affected across independent origins. The strongest cytonuclear signal involved the chloroplast, where dioecious branches exhibited relaxed selection in plastid coding sequences and nuclear-encoded plastid-targeted genes were enriched for episodic positive selection. Below, we consider how independent origins of dioecy, recurrent hybridization, organelle capture, and shifts in coding-sequence evolution together shaped the evolutionary history of the genus.

### Phylogenetic Relationships and Sexual-System Evolution

We resolved the deepest split among sampled *Amaranthus* species with genome-scale nuclear data and estimate that the sampled crown diversified approximately 2–5 Ma. The deepest divergence places *A. fimbriatus*, *A. tricolor*, and *A. viridis* as sister to all other sampled lineages. While *A. tricolor* and *A. viridis* have been recovered near the root of the genus in earlier phylogenies, the placement of *A. fimbriatus* has varied widely across studies (Raiyemo and Tranel 2023; Stetter and Schmid 2017; Viljoen et al. 2018; Waselkov et al. 2018; Xu et al. 2024). Our age estimate is broadly consistent with the ∼5 Ma estimate of Xu et al. (2024). Broader taxon sampling may refine relationships within this early-diverging group, but chromosome structure independently supports its separation from the rest of the genus. Sampled lineages outside this clade share a derived fusion of chromosomal segments homologous to *A. tricolor* chromosomes 1 and 2 (Raiyemo et al. 2025a,b). Under this topology, *A. viridis* and *A. fimbriatus* should retain the ancestral unfused configuration. Chromosome-level assemblies for these species would test whether this cytogenomic boundary marks the deepest split in *Amaranthus*.

The nuclear tree also confirmed that *Acnida* does not represent a single origin of dioecy. Species historically assigned to *Acnida* instead fell into two deeply separated clades, and ancestral-state reconstruction recovered at least two transitions from monoecy to dioecy (Mosyakin and Robertson 1996; Sauer 1955; Waselkov et al. 2018). Sex-determining regions in *A. palmeri* and *A. tuberculatus* corroborate this conclusion because they occur on different chromosomes, lack syntenic correspondence, and contain largely non-overlapping candidate gene sets (Bobadilla et al. 2023; Kreiner et al. 2025; Raiyemo et al. 2025a,b). The dioecious amaranths therefore evolved separate sexes through distinct genomic architectures rather than through inheritance of a single ancestral dioecious system.

The ancestral-state reconstruction revealed a likely return to monoecy in *A. pumilus* and left open a second possible loss in *A. spinosus* (Fig. 1). *A. pumilus* nested within Dioecious I, a clade otherwise composed of dioecious species, whereas *A. spinosus* fell as sister to *A. palmeri* and *A. watsonii* in Dioecious II, where the ancestral state remained ambiguous. *A. spinosus* is notable because male and female flowers occupy distinct regions of the same inflorescence rather than occurring in the mixed-sex glomerules typical of monoecious amaranths, a pattern that some authors interpreted as suggestive of a transition toward dioecy (Mosyakin and Robertson 1996; Waselkov et al. 2018). Whether this morphology reflects an incomplete transition toward dioecy or the residue of a return from dioecy, the placements of *A. pumilus* and *A. spinosus* raise broader questions about the conditions under which separate sexes are lost.

Reversals from dioecy to monoecy are documented in several angiosperm lineages (Barrett 2013), including at least seven independent transitions in *Momordica* (Schaefer and Renner 2010) and rapid breakdown of dioecy under mate limitation in *Mercurialis annua* (Cossard and Pannell 2021). Such reversals are often attributed to reproductive assurance, since obligate outcrossing leaves dioecious populations vulnerable when mates are scarce (Käfer et al. 2017; Renner and Ricklefs 1995). The ecology of *A. pumilus* fits this expectation, as it occupies ephemeral barrier-island habitats that are regularly disturbed and recolonized, where founding populations may consist of few individuals and self-compatible reproduction would be favored under Baker’s Law (Baker 1955; Pannell et al. 2015; Pannell and Barrett 1998). Hybridization provides another potential route to the breakdown of dioecy because divergent sex-determining systems can generate novel sexual phenotypes when brought together in the same genetic background (Alarcón-Bolaños et al. 2025; Yakimowski and Barrett 2016). Whether this mechanism contributed to the inferred reversion in *A. pumilus* cannot be determined from the present data, but the organelle-capture results place this lineage within a broader history of between-clade exchange.

The sex-determining systems in both dioecious clades retain characteristics consistent with an evolutionarily recent and incompletely canalized separation of sexes. In *A. tuberculatus*, Kreiner et al. (2025) identified a proto-sex chromosome in which recombination suppression is incomplete, Y haplotypes vary substantially in structure and gene content, and no single locus fully predicts phenotypic sex. They also documented inconstant sex expression, including individuals bearing flowers of both sexes and hermaphroditic flowers. Developmental studies of floral organogenesis in *A. palmeri* revealed that staminate flowers initially produce both male and female organ primordia before the gynoecium is aborted (Wu et al. 2023), indicating that the developmental program for female function has not been fully dismantled. These observations suggest that some capacity for combined sexual function persists in both clades and that the separation of sexes may remain subject to modification by genetic modifiers, environmental signals, or both.

### Reticulate Evolution and Organelle Capture

Hybridization shaped *Amaranthus* at more than one evolutionary scale, with ancient organelle capture crossing the boundary between dioecious clades and nuclear introgression concentrated mainly within those clades. Earlier plastid studies recovered part of the organellar pattern, especially the displacement of *A. palmeri* and *A. watsonii* from their nuclear relatives (Raiyemo and Tranel 2023; Waselkov et al. 2018), but incomplete lineage sorting remains a plausible explanation.

Coalescent simulations parameterized from genome-wide nuclear data and scaled for organellar inheritance placed the observed rearrangements far outside the ILS expectation. The results indicate that the Dioecious II ancestor of *A. palmeri* and *A. watsonii* acquired a plastome from an *A. pumilus-like* donor and a mitochondrion from an *A. acanthochiton-like* donor, both within Dioecious I. The mitochondrial tree also places *A. pumilus* outside its nuclear position within Dioecious I, consistent with a separate cytoplasmic exchange involving a donor from outside the dioecious clade. A third organelle capture displaced the *A. australis*/*A. cannabinus* wetland lineage from its nuclear position, although no sampled lineage could be identified as the donor.

The nuclear genome recorded a different history of hybridization. None of the nuclear introgression signals mapped to the events implicated by the organelle captures, consistent with the different population-genetic behavior of organellar and nuclear genomes. Plastid and mitochondrial genomes are typically uniparentally inherited, haploid, and effectively nonrecombining, giving the much smaller effective population sizes than nuclear loci (Rieseberg and Soltis 1991; Toews and Brelsford 2012; Tsitrone et al. 2003), so captured organelles can fix and persist while accompanying nuclear introgression is progressively eroded by recombination during backcrossing. Organelle capture without detectable accompanying nuclear introgression is widely documented across vascular plants (Stull et al. 2023).

The nuclear signals we did recover instead pointed to other episodes of gene flow. *A. watsonii* and *A. spinosus* showed strong evidence of localized nuclear introgression, consistent with more recent or less-eroded gene flow within Dioecious II. A broader reticulation signal connected the wetland lineage of *A. floridanus* and *A. tuberculatus* with sand-dwelling species within Dioecious I, while a deeper signal involving *A. fimbriatus* lacked the genomic structure expected from recent exchange and may reflect older introgression or gene flow from an unsampled lineage. These within-clade signals match experimental evidence for fertile crosses and herbicide-resistance introgression within dioecious clades (Gaines et al. 2012; Trucco et al. 2005a,b,c). The organellar and nuclear genomes therefore carry partly independent records of reticulate evolution in *Amaranthus*, with organelles preserving ancient between-clade exchange and nuclear data retaining stronger evidence of gene flow within clades.

The between-clade organelle captures occurred despite evidence from experimental crosses that hybridization between the two dioecious clades is substantially less successful than hybridization within either clade. Within-clade crosses have produced fertile offspring and documented introgression, whereas between-clade *A. palmeri* × *A. tuberculatus* crosses were recovered at low frequencies, and surviving hybrids frequently showed disrupted floral development, including neuter flowers, as well as chromosomal abnormalities (Gaines et al. 2012; Ju et al. 2025;

Oliveira et al. 2018; Trucco et al. 2005a,b,c, 2007). The two dioecious clades carry distinct sex-determining regions on different chromosomes with largely non-overlapping candidate gene sets (Bobadilla et al. 2023; Raiyemo et al. 2025a,b), so between-clade hybrids would bring divergent sex-determining architectures into the same genome. Comparable disruption of sex determination has been documented in other dioecious systems following admixture between divergent sexual systems (Alarcón-Bolaños et al. 2025; Yakimowski and Barrett 2016). A rare successful historical exchange followed by repeated backcrossing would nonetheless explain how an organelle genome crossed this boundary and persisted while nuclear donor ancestry fell below detection.

The inferred organelle captures localize to the Dioecious I sand-dwelling subclade, the same part of the phylogeny that contains *A. pumilus*, the only probable reversion to monoecy recovered in our ancestral-state reconstruction. The chloroplast capture implicates an *A. pumilus*-like lineage in the transfer of a Dioecious I plastome into the Dioecious II ancestor, while the mitochondrial placement of *A. pumilus* suggests possible additional cytoplasmic exchange involving a lineage outside Dioecious I, although support for that placement is limited. The lineage in which monoecy was regained therefore occurs within a part of the phylogeny marked by repeated cytoplasmic exchange and contact among lineages with contrasting sexual systems.

### Genome-Wide Molecular Responses to Dioecy

Dioecy in *Amaranthus* was associated with broad, largely lineage-specific shifts in coding-sequence evolution distributed across the nuclear genome rather than concentrated within characterized sex-determining regions. This pattern is consistent with the expectation that separating male and female function alters selection on many loci simultaneously. Genes involved in reproductive development, resource allocation, floral form, and gamete performance no longer contribute to fitness through both sex functions simultaneously (Charnov 1982; Lloyd 1982). As male and female fitness become decoupled, alleles constrained by shared fitness optima can evolve toward sex-specific values, sexually antagonistic variation can become exposed to sex-specific selection, and obligate outcrossing can alter the efficacy of selection through changes in mating system and effective population size (Barrett and Hough 2013; Charlesworth and Charlesworth 1978; Charlesworth 2018; Delph and Ashman 2006; Lande 1980; Lesaffre et al. 2025; Muyle et al. 2021).

Empirical studies of dioecious plants reveal diverse genomic responses to the evolution of separate sexes. In *Silene* subgenus *Silene*, autosomal and pseudoautosomal coding regions exhibited elevated positive selection on the branch where dioecy originated, even after excluding completely sex-linked genes (Zluvova et al. 2022). *Amaranthus* mirrored this pattern, with positive-selection signals concentrated on branches associated with the origins of dioecious clades but little overlap in the specific loci affected across independent origins (Fig. 5A). Dioecious *Trichosanthes* experienced accelerated evolution of male-biased genes through both positive selection and relaxed purifying selection (Zhao et al. 2024), whereas sex-biased genes in *Mercurialis* and *Leucadendron* exhibited no comparable increase in sequence-evolution rates, and faster expression evolution in *Leucadendron* appears to reflect ancestrally relaxed constraint rather than adaptation following the evolution of dioecy (Cossard and Pannell 2019; Scharmann et al. 2021). Together, these comparisons suggest that dioecy can reshape coding-sequence evolution, but that the targets and modes of response are strongly lineage dependent.

Only ten loci showed episodic positive selection on the stems marking the two independent origins of dioecy. The limited gene-level overlap parallels the distinct sex-determining architectures of *A. tuberculatus* and *A. palmeri*, which involve different chromosomes and largely non-syntenic candidate gene families (Bobadilla et al. 2023; Kreiner et al. 2025; Raiyemo et al. 2025a,b). Among the shared loci, CLV2, a CLAVATA-pathway receptor, carried signatures of positive selection on internal Dioecious I branches; CLAVATA/WUS meristem regulation contributes to sex-specific floral development in *Silene latifolia* (Kazama et al. 2022; Kobayashi et al. 2023), and CLAVATA2 and WUSCHEL were among sex-biased floral-development candidates in *A. tuberculatus* transcriptomes (Bobadilla et al. 2023). SEC8, required for pollen-tube growth (Cole et al. 2005), exhibited intensified selection in Dioecious I, consistent with stronger selection on male gametophytic performance after the shift to obligate outcrossing (Delph 2019). These examples point to plausible reproductive targets while emphasizing that most of the shared response remains functionally unresolved.

Most loci exhibiting altered selection dynamics lacked obvious reproductive annotations. This likely reflects both the breadth of the genomic response to dioecy and the conservative design of our analyses. We restricted our tests to single-copy orthogroups retained across the genus, excluding recent paralogs, lineage-specific genes, haplotype-specific sequence, and genes in regions where orthology assignment was uncertain. The analyses also targeted coding-sequence evolution and would not detect purely regulatory changes. The loci identified here therefore represent only a subset of the genomic response to dioecy. Rather than indicating that separate sexes affected a narrow set of functions, the scarcity of annotated reproductive genes suggests that much of the underlying response remains outside the conserved coding fraction of the genome sampled in this study.

### Selection at the Plastid-Nuclear Interface

Plastid coding sequences evolved under relaxed purifying selection on dioecious branches, and plastid-associated genes were the only enriched functional class among nuclear loci with altered selection dynamics. Together, these signals suggest that dioecy affected both the plastid genome and the nuclear genes that interact with it. Under maternal plastid inheritance, dioecy restricts plastid transmission to females, reducing plastid effective population size relative to the nuclear genome and weakening purifying selection (Birky et al. 1983; Postel and Touzet 2020). Maternal inheritance can also allow variants with male-specific fitness costs to persist, paralleling the mother’s curse framework developed for mitochondria (Frank and Hurst 1996; Gemmell et al. 2004; Havird et al. 2019). Whether plastids experience analogous sex-specific fitness effects in dioecious *Amaranthus* remains unknown, but relaxed purifying selection in plastid coding sequences is consistent with this possibility.

The co-occurrence of relaxed purifying selection in plastid coding sequences and altered selection on nuclear plastid-associated genes is consistent with cytonuclear evolution. Plastid function depends on interactions between plastid-encoded proteins and numerous imported nuclear proteins, creating opportunities for evolutionary change in one genome to influence selection in the other (Postel and Touzet 2020; Sloan 2015; Sloan et al. 2017; Weng et al. 2016). Chloroplast capture could also generate cytonuclear mismatch by introducing a divergent plastome into a novel nuclear background (Shahbazi et al. 2026), as in *A. palmeri* and *A. watsonii*. However, neither relaxed purifying selection in plastid coding sequences nor episodic selection on nuclear plastid-associated genes was restricted to lineages with captured plastomes. The strongest support occurred in

Dioecious I lineages retaining native plastomes, and selection on nuclear plastid-associated genes appeared on stem branches of both dioecious clades. The plastid-nuclear signal therefore aligns more closely with dioecy than with organelle capture alone, although the two processes may have interacted where both occurred.

Mitochondrial coding sequences underwent no comparable relaxation of purifying selection on dioecious branches. Plant mitochondrial genomes evolve much more slowly than plastid genomes at synonymous sites (Wolfe et al. 1987), and our mitochondrial alignment was smaller than the plastid alignment, reducing power to detect a parallel shift. Similar transmission asymmetries are expected for mitochondria and plastids in dioecious lineages because both are maternally inherited. Whether the absence of a mitochondrial signal reflects biological differences between the organellar genomes or limited power remains unresolved.

### Amaranthus as a Model for the Origin and Stability of Separate Sexes

Dioecy characterizes about six percent of angiosperm species despite having evolved hundreds to thousands of times (Renner 2014), a paradox likely explained in part by frequent reversions to combined sexual systems (Barrett 2013; Goldberg et al. 2017; Käfer et al. 2014, 2017; Wang et al. 2021). In *Amaranthus*, dioecy arose twice through non-homologous sex-determining regions on different chromosomes, produced largely non-overlapping coding-sequence responses across the two origins, and was lost at least once in *A. pumilus*. What selective conditions drove these independent transitions, and whether common ecological pressures produced parallel outcomes through different genetic mechanisms, remain open questions. Both dioecious clades retain young, homomorphic sex chromosomes with incomplete recombination suppression, fitting a broader pattern across animals and plants in which sex chromosomes frequently remain undifferentiated rather than progressing inevitably toward heteromorphy (Furman et al. 2020). In *A. tuberculatus*, Y haplotypes vary in structure and gene content, and inconstant sex expression has been reported (Kreiner et al. 2025). In *A. palmeri*, staminate flowers initiate both male and female organ primordia before gynoecium arrest, indicating that the developmental capacity for combined sexual function has not been irreversibly dismantled (Wu et al. 2023). This residual lability is the substrate on which selection for reproductive assurance can act, and experimental work has demonstrated that selection on comparable variation can rapidly restore functional hermaphroditism under mate limitation (Cossard and Pannell 2021). The barrier-island habitats of *A. pumilus*, subject to recurrent disturbance and colonization by small founding populations, provide conditions under which reproductive assurance would be strongly favored (Baker 1955; Pannell 2015). The organelle-capture results also situate *A. pumilus* within a part of the phylogeny shaped by repeated cytoplasmic exchange among lineages with contrasting sexual systems, raising the possibility that introgression introduced alleles that promoted reversion toward combined sexual function.

*A. palmeri* and *A. tuberculatus* are among the most damaging agricultural weeds globally; *A. palmeri* is now reported in at least 45 countries and is projected to spread into the temperate regions where *A. tuberculatus* is already established (Raiyemo et al. 2025a,b; Steckel 2007). Postzygotic barriers separate these species; experimental crosses yield hybrids only at low frequency and with disrupted floral development (Ju et al. 2025; Oliveira et al. 2018; Trucco et al. 2005a,b,c). The organelle captures nonetheless demonstrate that exchange between the two dioecious clades has occurred historically. Unlike plastids and mitochondria, which are maternally inherited through ovules, the ∼400 kb EPSPS-bearing extrachromosomal replicon conferring glyphosate resistance in *A. palmeri* replicates autonomously and is transmissible through pollen. This route has already carried the replicon naturally into *A. spinosus* and has been confirmed in experimental *A. palmeri* × *A. tuberculatus* crosses (Koo et al. 2018, 2023; Nandula et al. 2014). Historical organelle capture and contemporary transfer of adaptive genetic material indicate that reproductive barriers in *Amaranthus* are permeable enough for consequential exchange, even where hybridization is rare or developmentally disruptive. Population-genomic tests of admixture, paired with surveys of sex-expression variation, realized sex ratios, and mating-system parameters across species and environments, would connect the macroevolutionary transitions documented here to the microevolutionary processes that produce, maintain, and reverse separate sexes.

## Acknowledgments

We thank Sandra Hoffberg for plant cultivation, Jasmina Dzurlic for DNA extraction and sample preparation, and Spencer C. H. Barrett for comments on the manuscript.

## Data Availability

Sequencing reads for the 14 newly generated whole-genome accessions have been deposited in the NCBI Short Read Archive under BioProject PRJNA1476526, and supporting data, analysis files, and code will be made available in the Dryad Digital Repository upon publication.

## Supplementary Material

Supplementary material includes supplementary tables S1–S10 and is available as a supplementary file accompanying this manuscript.

