## Supplementary Tables for "Organelle Capture, Lineage-Specific Genomic Responses, and the Lability of Dioecy in *Amaranthus*"

### Supplementary Material

Supplementary Table S1. Sample information and accessions for all taxa included in the phylogenomic analyses. Tip name is the label used on the species tree (Fig. 1) and cytoplasmic phylogenies (Fig. 2). Sequence accession reports public sequence/archive or reference identifiers where available. GRIN accession reports USDA plant-material identifiers for samples sequenced in this study. Closest reference indicates the reference genome used for consensus-sequence reconstruction where applicable.

| Tip name | Data type | Source | Sequence accession | GRIN accession | Closest reference |
|---|---|---|---:|---:|---|
| *A. fimbriatus* USDA 2021 | WGS | This study | SRR39067046 | PI 662285 | *A. tricolor* |
| *A. viridis* SRA1 | WGS | SRA | SRR7121710 | — | *A. tricolor* |
| *A. tricolor* NCBI ref | Reference genome | NCBI | ASM2621246v1 | — | — |
| *A. tricolor* SRA1 | WGS | SRA | SRR17777275 | — | *A. tricolor* |
| *A. tricolor* SRA2 | WGS | SRA | SRR21968681 | — | *A. tricolor* |
| *A. retroflexus* CoGe ref | Reference genome | CoGe | 68315 | — | — |
| *A. retroflexus* USDA 2021 | WGS | This study | SRR39067045 | PI 607447 | *A. retroflexus* |
| *A. caudatus* USDA 2021 | WGS | This study | SRR39067040 | Ames 15145 | *A. hypochondriacus* |
| *A. cruentus* USDA 2021 | WGS | This study | SRR39067039 | PI 576461 | *A. hypochondriacus* |
| *A. hybridus* CoGe ref | Reference genome | CoGe | 68316 | — | — |
| *A. hybridus* SRA1 | WGS | SRA | SRR12075659 | — | *A. hybridus* |
| *A. hybridus* SRA2 | WGS | SRA | SRR14055740 | — | *A. hybridus* |
| *A. hypochondriacus* NCBI ref | Reference genome | NCBI | GCA_977020195.1 | — | — |
| *A. hypochondriacus* SRA1 | WGS | SRA | SRR2130053 | — | *A. hypochondriacus* |
| *A. hypochondriacus* SRA2 | WGS | SRA | SRR2130055 | — | *A. hypochondriacus* |
| *A. spinosus* USDA 2021 | WGS | This study | SRR39067038 | PI 619234 | *A. palmeri* |
| *A. spinosus* SRA1 | WGS | SRA | SRR7121582 | — | *A. palmeri* |
| *A. palmeri* CoGe ref | Reference genome | CoGe | 68302 | — | — |
| *A. palmeri* F USDA P04 | WGS | This study | SRR39067036 | — | *A. palmeri* |
| *A. palmeri* M USDA P12 | WGS | This study | SRR39067035 | — | *A. palmeri* |
| *A. watsonii* F SRA1 | WGS | SRA | SRR19158638 | — | *A. palmeri* |
| *A. watsonii* M SRA2 | WGS | SRA | SRR19158646 | — | *A. palmeri* |
| *A. cannabinus* F SRA1 | WGS | SRA | SRR19158643 | — | *A. tuberculatus* |

| Tip name | Data type | Source | Sequence accession | GRIN accession | Closest reference |
|---|---|---|---|---|---|
| *A. cannabinus* M USDA 2021 | WGS | This study | SRR39067037 | PI 641041 | *A. tuberculatus* |
| *A. cannabinus* M SRA2 | WGS | SRA | SRR19158642 | — | *A. tuberculatus* |
| *A. australis* M SRA1 | WGS | SRA | SRR19158644 | — | *A. tuberculatus* |
| *A. floridanus* M SRA1 | WGS | SRA | SRR19158641 | — | *A. tuberculatus* |
| *A. tuberculatus* CoGe ref | Reference genome | CoGe | 69059 | — | — |
| *A. tuberculatus* F USDA T02 | WGS | This study | SRR39067034 | — | *A. tuberculatus* |
| *A. tuberculatus* M USDA SD008 | WGS | This study | SRR39067033 | — | *A. tuberculatus* |
| *A. acanthochiton* F SRA1 | WGS | SRA | SRR19158648 | — | *A. tuberculatus* |
| *A. acanthochiton* M SRA2 | WGS | SRA | SRR19158647 | — | *A. tuberculatus* |
| *A. greggii* USDA AGF1 | WGS | This study | SRR39067044 | PI 667170 | *A. tuberculatus* |
| *A. greggii* USDA AGM11 | WGS | This study | SRR39067043 | PI 667170 | *A. tuberculatus* |
| *A. arenicola* SRA1 | WGS | SRA | SRR19158645 | — | *A. tuberculatus* |
| *A. pumilus* DS002 | WGS | This study | SRR39067042 | — | *A. tuberculatus* |
| *A. pumilus* EP001 | WGS | This study | SRR39067041 | — | *A. tuberculatus* |
| *Alt. philoxeroides* NCBI ref | Transcriptome | NCBI | — | — | — |
| *D. amaranthoides* NCBI ref | Transcriptome | NCBI | — | — | — |
| *H. glauca* NCBI ref | Transcriptome | NCBI | — | — | — |

Supplementary Table S2. Nuclear sequence recovery and alignment summary for the *Amaranthus* single-copy coding dataset. Outgroup transcriptome rows report locus contribution to the retained orthogroup set.

| Category | Metric | Value | Notes |
|---|---|---:|---|
| Nuclear orthogroups | single-copy coding alignments retained | 4,329 | After orthogroup filtering, isoform resolution, codon-aware alignment, and trimming. |
| Nuclear orthogroups | total aligned columns | 5,889,951 | Sum across retained nucleotide CDS alignments. |
| Nuclear orthogroups | mean alignment length (bp) | 1,361 | Median length was 1,164 bp. |
| Nuclear orthogroups | mean taxa per alignment | 21.57 | Maximum observed taxon count was 22. |
| Nuclear orthogroups | mean occupancy (%) | 98.02 | Occupancy is relative to the maximum observed taxon count. |
| Whole-genome accessions | median nuclear depth (x) | 13.15 | Computed across 29 ingroup WGS accessions with samtools-stats coverage values; range 8.07-45.4x. |
| Whole-genome accessions | median orthogroups recovered | 4,329 | Computed across 31 ingroup accessions with consensus-recovery results; range 4,325-4,329. |
| Outgroup transcriptomes | *Alternanthera philoxeroides* | 3,942 loci (91.1%) | Mean recovered CDS length 1,072 bp. |
| Outgroup transcriptomes | *Deeringia amaranthoides* | 3,789 loci (87.5%) | Mean recovered CDS length 1,168 bp. |
| Outgroup transcriptomes | *Hermbstaedtia glauca* | 3,925 loci (90.7%) | Mean recovered CDS length 1,158 bp. |

Supplementary Table S3. Posterior parameter estimates from the completed BPP A00 species-tree analysis on filtered nuclear single-copy orthogroup alignments under the rooted *Amaranthus* ASTRAL species tree. Estimates are from the completed A00 chain.

| Node | Lineage | Type | θ (95% HPD) | θ ESS | τ (95% HPD) | τ ESS |
|---|---|---|---|---|---|---|
| 1 | *A. tricolor* | tip | 2.3e-4 (2.1e-4–2.4e-4) | 3,062 | — | — |
| 2 | *A. viridis* | tip | 1.7e-4 (1.5e-4–1.9e-4) | 5,794 | — | — |
| 3 | *A. fimbriatus* | tip | 5.8e-4 (5.3e-4–6.3e-4) | 12,743 | — | — |
| 4 | *A. hybridus* | tip | 6.5e-5 (5.7e-5–7.4e-5) | 8,049 | — | — |
| 5 | *A. hypochondriacus* | tip | 4.8e-5 (4.2e-5–5.4e-5) | 10,523 | — | — |
| 6 | *A. cruentus* | tip | 5.1e-5 (4.5e-5–5.9e-5) | 8,609 | — | — |
| 7 | *A. caudatus* | tip | 5.7e-5 (4.9e-5–6.5e-5) | 9,563 | — | — |
| 8 | *A. retroflexus* | tip | 3.6e-4 (3.4e-4–3.8e-4) | 9,034 | — | — |
| 9 | *A. palmeri* | tip | 0.0199 (0.0190–0.0207) | 2,796 | — | — |
| 10 | *A. watsonii* | tip | 0.0082 (0.0078–0.0086) | 951 | — | — |
| 11 | *A. spinosus* | tip | 6.1e-4 (5.8e-4–6.4e-4) | 753 | — | — |
| 12 | *A. floridanus* | tip | 0.0032 (0.0030–0.0034) | 11,469 | — | — |
| 13 | *A. tuberculatus* | tip | 0.0131 (0.0126–0.0136) | 6,723 | — | — |
| 14 | *A. cannabinus* | tip | 0.0034 (0.0033–0.0035) | 6,263 | — | — |
| 15 | *A. australis* | tip | 0.0015 (0.0014–0.0016) | 15,576 | — | — |
| 16 | *A. pumilus* | tip | 3.4e-4 (3.1e-4–3.6e-4) | 7,307 | — | — |
| 17 | *A. acanthochiton* | tip | 0.0066 (0.0063–0.0068) | 5,439 | — | — |
| 18 | *A. greggii* | tip | 0.0016 (0.0015–0.0017) | 7,065 | — | — |
| 19 | *A. arenicola* | tip | 0.0043 (0.0040–0.0047) | 16,804 | — | — |
| 20 | *Amaranthus* crown (all 19 ingroup taxa) | internal | 0.0150 (0.0144–0.0156) | 1,364 | 0.0140 (0.0137–0.0143) | 1,251 |
| 21 | Early-diverging clade (tricolor + viridis + fimbriatus) | internal | 0.0171 (0.0156–0.0186) | 7,097 | 0.0070 (0.0068–0.0073) | 3,532 |
| 22 | tricolor + viridis | internal | 9.5e-4 (6.8e-4–0.0012) | 4,832 | 0.0056 (0.0054–0.0058) | 3,241 |
| 23 | Hybridus/grain + Dioecious I + Dioecious II (16 taxa) | internal | 0.0026 (3.1e-4–0.0071) | 6,054 | 0.0140 (0.0137–0.0143) | 1,249 |
| 24 | Hybridus/grain + Dioecious II | internal | 0.0064 (0.0058–0.0070) | 3,778 | 0.0106 (0.0103–0.0108) | 1,965 |

| Node | Lineage | Type | θ (95% HPD) | θ ESS | τ (95% HPD) | τ ESS |
|---|---|---|---|---|---|---|
| 25 | Hybridus/grain crown (5 taxa) | internal | 0.0066 (0.0062–0.0069) | 8,287 | 0.0016 (0.0015–0.0016) | 7,763 |
| 26 | hybridus + hypochondriacus + cruentus + caudatus | internal | 0.0016 (0.0014–0.0018) | 14,515 | 0.0008 (0.0007–0.0008) | 8,100 |
| 27 | hybridus + hypochondriacus | internal | 0.0018 (0.0017–0.0019) | 6,956 | 2.4e-5 (2.1e-5–2.7e-5) | 6,592 |
| 28 | cruentus + caudatus | internal | 7.6e-4 (7.0e-4–8.2e-4) | 16,640 | 7.2e-5 (6.2e-5–8.1e-5) | 5,450 |
| 29 | Dioecious II (palmeri + watsonii + spinosus) | internal | 0.0124 (0.0120–0.0129) | 220 | 0.0023 (0.0023–0.0024) | 91 |
| 30 | palmeri + watsonii | internal | 0.0043 (2.8e-4–0.0103) | 3,838 | 0.0023 (0.0023–0.0024) | 93 |
| 31 | Dioecious I (8 taxa) | internal | 0.0167 (0.0161–0.0173) | 3,526 | 0.0050 (0.0049–0.0051) | 2,096 |
| 32 | Wetland subclade (floridanus + tuberculatus + cannabinus + australis) | internal | 0.0174 (0.0138–0.0210) | 6,321 | 0.0044 (0.0042–0.0045) | 2,419 |
| 33 | floridanus + tuberculatus | internal | 0.0217 (0.0196–0.0238) | 10,666 | 0.0027 (0.0025–0.0028) | 3,664 |
| 34 | cannabinus + australis | internal | 0.0054 (0.0050–0.0057) | 8,966 | 0.0019 (0.0018–0.0020) | 4,242 |
| 35 | Sand-dwelling subclade (pumilus + acanthochiton + greggii + arenicola) | internal | 0.0093 (0.0060–0.0127) | 4,949 | 0.0045 (0.0043–0.0047) | 2,761 |
| 36 | acanthochiton + greggii + arenicola | internal | 0.0086 (0.0066–0.0106) | 6,231 | 0.0037 (0.0036–0.0039) | 3,168 |
| 37 | greggii + arenicola | internal | 0.0147 (0.0133–0.0161) | 11,906 | 0.0016 (0.0015–0.0017) | 3,968 |

Supplementary Table S4. Variant filtering and substitution-composition summary for the SNP dataset analyzed with Dsuite. The final dataset contained 30 *Amaranthus* ingroup accessions plus *Alternanthera philoxeroides* as outgroup and was aligned to the *A. tricolor* reference.

| Section | Metric | Value | Notes |
|---|---|---:|---|
| Variant filtering | Biallelic SNPs QUAL25 SnpGap5 genotype QC missingness 20% | 15,560,632 | Per-contig filters: QUAL>=25; SnpGap 5; mask DP<8 OR GQ<30 OR SP>60; F_MISSING<=0.2. |
| Variant filtering | Plus depth ceiling 2x mean | 11,099,793 | Post-hoc: INFO/DP <= 1779 (= 2x mean 889.9); MAC>=2 jointly applied. |
| Variant filtering | Final sites into Dsuite | 11,099,793 | Final genome-wide biallelic SNP set used for D / f4 / fbranch. |
| Variant filtering | Ts/Tv ratio | 1.46 | Transitions/transversions on final SNP set (6587642 ts / 4512151 tv). |
| Variant filtering | N samples | 31 | Includes *Amaranthus* ingroup + Alternanthera outgroup. |
| Substitution composition | Transition SNPs | 6,587,642 | Summed from the final SNP set substitution-type table. |
| Substitution composition | Transversion SNPs | 4,512,151 | Summed from the final SNP set substitution-type table. |

Supplementary Table S5. Posterior summaries from BPP MSci quartet analyses retained for manuscript interpretation. The introgression fraction is reported as 1 - phi with 95% HPD interval in parentheses. Non-converged, incomplete, and exploratory post-processed bidirectional model fits were excluded.

| Quartet | Model | Chain | Direction | 1 − φ | ESS | ln L | R-hat |
|---|---|---|---|---|---|---|---|
| *viridis–fimbriatus* | FV (fimbriatus → viridis) | 1 | fimb → vir | 0.0012 (0.0000–0.0033) | 14,595 | -2,062,672 | 1.000 |
| *viridis–fimbriatus* | FV (fimbriatus → viridis) | 2 | fimb → vir | 0.0012 (0.0000–0.0033) | 17,215 | -2,062,679 | — |
| *watsonii–spinosus* | Base tree (no admixture) | 1 | — | — | — | -2,248,827 | — |
| *watsonii–spinosus* | Base tree (no admixture) | 2 | — | — | — | -2,248,869 | — |
| *watsonii–spinosus* | SW (spinosus → watsonii) | 1 | spin → wat | 0.1429 (0.1304–0.1557) | 1,802 | -2,246,866 | 1.000 |
| *watsonii–spinosus* | SW (spinosus → watsonii) | 2 | spin → wat | 0.1428 (0.1304–0.1557) | 2,814 | -2,246,898 | — |
| *watsonii–spinosus* | WS (watsonii → spinosus) | 1 | wat → spin | 0.4695 (0.4379–0.5011) | 4,284 | -2,246,859 | 1.000 |
| *watsonii–spinosus* | WS (watsonii → spinosus) | 2 | wat → spin | 0.4700 (0.4384–0.5017) | 3,642 | -2,246,857 | — |

Supplementary Table S6. Manually curated dioecy-relevant Gene Ontology biological-process terms grouped into broad categories for descriptive functional summaries. The curated list was used for screening candidate genes and was not treated as a formal enrichment test.

| Category | GO ID | GO Term | Relevance |
|---|---|---|---|
| Reproductive/floral development | GO:0022414 | Reproductive process | Broad reproduction |
| | GO:0000003 | Reproduction | Broadest reproductive bin |
| | GO:0019953 | Sexual reproduction | Sexual reproduction specifically |
| | GO:0003006 | Developmental process involved in reproduction | Reproductive development |
| | GO:0048608 | Reproductive structure development | Reproductive organs/structures |
| | GO:0090567 | Reproductive shoot system development | Flowering/reproductive shoot context |
| | GO:0009908 | Flower development | Central floral-development parent |
| | GO:0048437 | Floral organ development | Parent of stamen, carpel, petal, sepal development |
| | GO:0048444 | Floral organ morphogenesis | Organ shape/form |
| | GO:0048449 | Floral organ formation | Initiation/formation of floral organs |
| | GO:0048450 | Floral organ structural organization | Organization of floral organs |
| | GO:0048497 | Maintenance of floral organ identity | Organ identity maintenance |
| | GO:0048833 | Specification of floral organ number | Organ-number patterning |
| | GO:0009909 | Regulation of flower development | Regulation of flower development |
| | GO:0009911 | Positive regulation of flower development | Enhanced/promoted development |
| Male organ/fertility | GO:0048466 | Androecium development | Male floral whorl development; useful for genes affecting the collective stamen-bearing organ system. |
| | GO:0048443 | Stamen development | Core male floral organ development term; directly relevant to male-function evolution. |
| | GO:0048448 | Stamen morphogenesis | Shape and structural development of stamens; relevant to altered male organ morphology. |
| | GO:0048455 | Stamen formation | Initiation/formation of male floral organs; relevant to stamen suppression or loss. |
| | GO:0010097 | Specification of stamen identity | Male floral organ identity specification; directly relevant to shifts in stamen fate or suppression. |
| | GO:0048653 | Anther development | Core anther development term; directly relevant to pollen-bearing organ function. |
| | GO:0048654 | Anther morphogenesis | Anther shape and tissue organization; relevant to male organ function. |

| Category | GO ID | GO Term | Relevance |
|---|---|---|---|
| | GO:0009901 | Anther dehiscence | Anther opening and pollen release; relevant to functional male fertility. |
| | GO:0120194 | Regulation of anther dehiscence | Regulation of anther opening; relevant to pollen release and male fertility. |
| | GO:0120195 | Positive regulation of anther dehiscence | Promotion of anther opening; relevant to pollen release and male fertility. |
| | GO:0120196 | Negative regulation of anther dehiscence | Suppression of anther opening; relevant when pollen is produced but not released. |
| | GO:0048658 | Anther wall tapetum development | Tapetum development; strong candidate process for male sterility or pollen abortion. |
| | GO:0048656 | Anther wall tapetum formation | Tapetal tissue formation; relevant to anther development and pollen fertility. |
| | GO:0048655 | Anther wall tapetum morphogenesis | Tapetal tissue morphogenesis; relevant to anther tissue organization and pollen support. |
| | GO:0048657 | Anther wall tapetum cell differentiation | Tapetal differentiation; relevant to pollen nourishment, maturation, and male sterility. |
| | GO:0010234 | Anther wall tapetum cell fate specification | Tapetal cell fate specification; important because tapetum defects commonly disrupt pollen fertility. |
| | GO:0009556 | Microsporogenesis | Male meiosis and microspore formation; directly relevant to male gamete production. |
| | GO:0055046 | Microgametogenesis | Male gametophyte development after microspore formation; relevant to pollen function. |
| | GO:0010480 | Microsporocyte differentiation | Differentiation of male meiotic cells; relevant to initiation of male gametophyte production. |
| | GO:0009555 | Pollen development | Core male gametophyte development term; directly relevant to male fertility. |
| | GO:0010152 | Pollen maturation | Pollen maturation; relevant to production of functional male gametophytes. |
| | GO:0048235 | Pollen sperm cell differentiation | Male gamete differentiation within pollen; relevant to functional male fertility. |
| | GO:0010208 | Pollen wall assembly | Pollen wall construction; relevant to viable pollen formation and male fertility. |
| | GO:0010584 | Pollen exine formation | Pollen exine formation; relevant to pollen protection, recognition, and viability. |

| Category | GO ID | GO Term | Relevance |
|---|---|---|---|
| | GO:0160030 | Pollen intine formation | Pollen intine formation; relevant to pollen structure and germination competence. |
| | GO:0062075 | Pollen aperture formation | Pollen aperture formation; relevant to pollen germination and tube emergence. |
| | GO:0080110 | Sporopollenin biosynthetic process | Sporopollenin biosynthesis; key process for exine formation and viable pollen. |
| Female organ/fertility | GO:0048467 | Gynoecium development | Female floral whorl development; relevant to carpel-bearing organ system evolution. |
| | GO:0048440 | Carpel development | Core female floral organ development term; directly relevant to female-function evolution. |
| | GO:0048445 | Carpel morphogenesis | Carpel shape and structural development; relevant to altered female organ morphology. |
| | GO:0048462 | Carpel formation | Initiation/formation of carpels; relevant to female organ development or suppression. |
| | GO:0048463 | Carpel structural organization | Carpel structural organization; relevant to functional gynoecium formation. |
| | GO:0010094 | Specification of carpel identity | Female floral organ identity specification; directly relevant to shifts in carpel fate. |
| | GO:0035670 | Plant-type ovary development | Ovary development; relevant to ovule-bearing female reproductive structures. |
| | GO:0080126 | Ovary septum development | Ovary development; relevant to ovule-bearing female reproductive structures. |
| | GO:0048481 | Plant ovule development | Core ovule development term; directly relevant to female fertility. |
| | GO:0048482 | Plant ovule morphogenesis | Ovule shape and structural development; relevant to female fertility and seed initiation. |
| | GO:0010622 | Specification of ovule identity | Ovule identity specification; relevant to initiation of female reproductive units. |
| | GO:0080060 | Integument development | Ovule integument development; relevant to ovule structure and seed precursor formation. |
| | GO:0048480 | Stigma development | Stigma development; relevant to pollen receipt and female reproductive function. |
| | GO:0048479 | Style development | Style development; relevant to pollen tube passage through female tissue. |

| Category | GO ID | GO Term | Relevance |
|---|---|---|---|
| | GO:0010500 | Transmitting tissue development | Pollen tube transmitting tissue development; relevant to female-side support of fertilization. |
| | GO:0009554 | Megasporogenesis | Female meiosis and megaspore formation; directly relevant to female gamete production. |
| | GO:0009561 | Megagametogenesis | Female gametophyte development; relevant to embryo sac formation and female fertility. |
| | GO:0009553 | Embryo sac development | Core female gametophyte development term; directly relevant to female fertility. |
| | GO:0048314 | Embryo sac morphogenesis | Embryo sac morphology; relevant to female gametophyte structure and function. |
| | GO:0009558 | Embryo sac cellularization | Cellularization of the embryo sac; relevant to normal female gametophyte development. |
| | GO:0009559 | Embryo sac central cell differentiation | Central-cell differentiation in the female gametophyte; relevant to embryo sac function. |
| | GO:0009560 | Embryo sac egg cell differentiation | Egg-cell differentiation in the female gametophyte; directly relevant to female fertility. |
| | GO:0045691 | Regulation of embryo sac central cell differentiation | Central-cell differentiation in the female gametophyte; relevant to embryo sac function. |
| | GO:0045692 | Negative regulation of embryo sac central cell differentiation | Suppression of central-cell differentiation; relevant to embryo sac defects and female fertility loss. |
| | GO:0045693 | Positive regulation of embryo sac central cell differentiation | Promotion of central-cell differentiation; relevant to functional embryo sac development. |
| | GO:0045694 | Regulation of embryo sac egg cell differentiation | Egg-cell differentiation in the female gametophyte; directly relevant to female fertility. |
| | GO:0045695 | Negative regulation of embryo sac egg cell differentiation | Suppression of egg-cell differentiation; relevant to female gametophyte defects. |
| | GO:0045696 | Positive regulation of embryo sac egg cell differentiation | Promotion of egg-cell differentiation; relevant to functional female gamete formation. |
| | GO:0009563 | Synergid differentiation | Synergid-cell differentiation; relevant to pollen tube attraction and fertilization. |
| | GO:0045697 | Regulation of synergid differentiation | Synergid-cell differentiation; relevant to pollen tube attraction and fertilization. |
| | GO:0045698 | Negative regulation of synergid differentiation | Suppression of synergid differentiation; relevant to female gametophyte defects. |

| Category | GO ID | GO Term | Relevance |
|---|---|---|---|
| | GO:0045699 | Positive regulation of synergid differentiation | Promotion of synergid differentiation; relevant to pollen tube reception. |
| | GO:0010198 | Synergid death | Programmed death of synergid cells; relevant to pollen tube reception and fertilization. |
| | GO:0055045 | Antipodal cell degeneration | Degeneration of antipodal cells; a female-gametophyte cell-death process. |
| Pollination/fertilization | GO:0009856 | Pollination | Pollen transfer and early reproductive interaction; directly relevant to realized mating success. |
| | GO:0009875 | Pollen-pistil interaction | Interaction between male gametophyte and female tissues; relevant to compatibility and fertility. |
| | GO:0048544 | Recognition of pollen | Pollen recognition by female tissues; relevant to compatibility and mate filtering. |
| | GO:0060321 | Acceptance of pollen | Acceptance of pollen; relevant to successful pollen-pistil compatibility. |
| | GO:0060320 | Rejection of self pollen | Self-pollen rejection; relevant to mating-system evolution and compatibility barriers. |
| | GO:1990109 | Rejection of pollen from other species | Heterospecific pollen rejection; relevant to reproductive isolation and pollen-pistil compatibility. |
| | GO:0009859 | Pollen hydration | Pollen hydration on the stigma; necessary for germination and male gametophyte function. |
| | GO:0009846 | Pollen germination | Pollen germination; required for pollen tube formation and male fertility. |
| | GO:0048868 | Pollen tube development | Development of the pollen tube; relevant to male gametophyte performance. |
| | GO:0009860 | Pollen tube growth | Pollen tube elongation; directly relevant to fertilization success. |
| | GO:0080092 | Regulation of pollen tube growth | Control of pollen tube elongation; relevant to successful delivery of sperm cells. |
| | GO:0010183 | Pollen tube guidance | Directed pollen tube growth toward the female gametophyte; central to fertilization success. |
| | GO:0160068 | Negative regulation of pollen tube guidance | Suppression of pollen tube guidance; relevant to failed fertilization after pollination. |
| | GO:0009865 | Pollen tube adhesion | Adhesion of pollen tube to female tissues; relevant to pollen tube passage and guidance. |

| Category | GO ID | GO Term | Relevance |
|---|---|---|---|
| | GO:0009566 | Fertilization | Union of gametes; endpoint of male and female reproductive function. |
| | GO:0009567 | Double fertilization forming a zygote and endosperm | Seed-plant-specific fertilization process forming embryo and endosperm. |
| | GO:0080155 | Regulation of double fertilization forming a zygote and endosperm | Regulation of double fertilization; relevant to successful seed initiation. |
| | GO:0061936 | Fusion of sperm to egg plasma membrane involved in double fertilization forming a zygote and endosperm | Gamete membrane fusion during double fertilization; directly relevant to fertilization success. |
| Sex-organ suppression/abortion | GO:0012501 | Programmed cell death | Cell death process; possible proxy for organ abortion or sex-specific tissue degeneration. |
| | GO:0043067 | Regulation of programmed cell death | Control of cell death; relevant to developmental suppression or degeneration of sex organs. |
| | GO:0043068 | Positive regulation of programmed cell death | Promotion of cell death; relevant to sex-organ degeneration or abortion-like processes. |
| | GO:0043069 | Negative regulation of programmed cell death | Suppression of cell death; relevant to persistence or rescue of reproductive tissues. |
| | GO:0010623 | Programmed cell death involved in cell development | Developmentally regulated cell death; relevant to organ sculpting, degeneration, or abortion. |
| | GO:0010421 | Hydrogen peroxide-mediated programmed cell death | ROS-mediated programmed cell death; possible mechanism for floral organ degeneration. |
| | GO:1901298 | Regulation of hydrogen peroxide-mediated programmed cell death | ROS-mediated programmed cell death; possible mechanism for floral organ degeneration. |
| | GO:1901299 | Negative regulation of hydrogen peroxide-mediated programmed cell death | Regulation of ROS-mediated cell death; possible mechanism for preventing organ degeneration. |
| | GO:1901300 | Positive regulation of hydrogen peroxide-mediated programmed cell death | Promotion of ROS-mediated cell death; possible mechanism for organ abortion or degeneration. |
| | GO:0090693 | Plant organ senescence | Organ senescence; relevant to reproductive tissue aging or degeneration. |
| | GO:0080187 | Floral organ senescence | Senescence of floral organs; possible proxy for sex-organ degeneration. |
| | GO:0010227 | Floral organ abscission | Floral organ shedding; possible proxy for organ loss. |
| | GO:0060863 | Regulation of floral organ abscission by signal transduction | Regulation of floral organ shedding by signaling; relevant to organ loss or retention. |

| Category | GO ID | GO Term | Relevance |
|---|---|---|---|
| | GO:0060861 | Positive regulation of floral organ abscission | Promotion of floral organ shedding; possible proxy for organ loss. |
| | GO:0060862 | Negative regulation of floral organ abscission | Suppression of floral organ shedding; relevant to persistence of reproductive organs. |
| | GO:0009910 | Negative regulation of flower development | Potential proxy for floral organ suppression; interpret with organ-specific context. |
| | GO:2000241 | Regulation of reproductive process | Broad regulation of reproductive function; interpret with more specific annotations. |
| | GO:2000242 | Negative regulation of reproductive process | Broad proxy for suppression of reproductive function; interpret cautiously. |
| | GO:2000243 | Positive regulation of reproductive process | Broad reproductive promotion term; useful for regulatory context. |
| Meiosis/recombination | GO:0051321 | Meiotic cell cycle | Broad meiosis term; relevant to gamete production and meiotic chromosome behavior. |
| | GO:1903046 | Meiotic cell cycle process | Broad meiosis term; relevant to gamete production and meiotic chromosome behavior. |
| | GO:0007127 | Meiosis I | First meiotic division; relevant to homolog segregation and gamete formation. |
| | GO:0061982 | Meiosis I cell cycle process | First meiotic division; relevant to homolog segregation and gamete formation. |
| | GO:0140013 | Meiotic nuclear division | Nuclear division during meiosis; relevant to gametogenesis. |
| | GO:0045132 | Meiotic chromosome segregation | Meiotic chromosome segregation; relevant to gamete viability and sex chromosome behavior. |
| | GO:0051307 | Meiotic chromosome separation | Separation of chromosomes during meiosis; relevant to gamete formation. |
| | GO:0007129 | Homologous chromosome pairing at meiosis | Homolog pairing during meiosis; relevant to recombination and chromosome segregation. |
| | GO:0070193 | Synaptonemal complex organization | Synapsis machinery; relevant to homolog pairing and recombination. |
| | GO:0007130 | Synaptonemal complex assembly | Synapsis machinery; relevant to homolog pairing and recombination. |
| | GO:0006310 | DNA recombination | Broad recombination term; relevant but not meiosis-specific. |

| Category | GO ID | GO Term | Relevance |
|---|---|---|---|
| | GO:0035825 | Homologous recombination | Homology-directed recombination; relevant to DNA repair and meiotic exchange. |
| | GO:0140527 | Reciprocal homologous recombination | Reciprocal exchange between homologous sequences; relevant to crossover-like events. |
| | GO:0007131 | Reciprocal meiotic recombination | Central crossover/recombination term for meiotic exchange. |
| | GO:0051026 | Chiasma assembly | Cytological manifestation of meiotic crossover; relevant to recombination and segregation. |
| | GO:0010780 | Meiotic DNA double-strand break formation involved in reciprocal meiotic recombination | Meiotic DSB initiation; upstream step in meiotic recombination. |
| | GO:0010705 | Meiotic DNA double-strand break processing involved in reciprocal meiotic recombination | Processing of meiotic DSBs; relevant to repair pathway choice and recombination. |
| | GO:1905261 | Regulation of meiotic DNA double-strand break formation involved in reciprocal meiotic recombination | Regulation of meiotic DSB initiation; relevant to crossover number and placement. |
| | GO:0010772 | Meiotic DNA recombinase assembly involved in reciprocal meiotic recombination | Assembly of meiotic recombinase machinery; relevant to strand invasion and recombination. |
| | GO:0010774 | Meiotic strand invasion involved in reciprocal meiotic recombination | Strand invasion during meiotic recombination; core step in homologous exchange. |
| | GO:0010777 | Meiotic mismatch repair involved in reciprocal meiotic recombination | Mismatch repair during meiotic recombination; relevant to heteroduplex resolution/gene conversion. |
| | GO:0010778 | Meiotic DNA repair synthesis involved in reciprocal meiotic recombination | DNA synthesis during meiotic repair; relevant to recombination intermediates. |
| | GO:0000709 | Meiotic joint molecule formation | Formation of meiotic recombination intermediates; relevant to crossover/noncrossover outcomes. |
| | GO:0000712 | Resolution of meiotic recombination intermediates | Resolution of meiotic recombination intermediates; relevant to crossover completion. |
| | GO:1990918 | Double-strand break repair involved in meiotic recombination | Meiotic DSB repair; links DNA repair machinery to recombination outcomes. |
| | GO:0010520 | Regulation of reciprocal meiotic recombination | Control of meiotic recombination; relevant to crossover frequency and sex-linked regions. |
| | GO:0045128 | Negative regulation of reciprocal meiotic recombination | Suppression of meiotic recombination; relevant to recombination reduction and sex chromosome evolution. |
| | GO:0010845 | Positive regulation of reciprocal meiotic recombination | Promotion of meiotic recombination; relevant to crossover control. |

| Category | GO ID | GO Term | Relevance |
|---|---|---|---|
| | GO:0006311 | Meiotic gene conversion | Non-reciprocal sequence transfer during meiosis; relevant to recombination and repair outcomes. |
| Hormone/signaling | GO:0007165 | Signal transduction | Very broad signaling term; useful only as a secondary regulatory category. |
| | GO:0009755 | Hormone-mediated signaling pathway | Plant hormone signaling; hormones can mediate floral organ development, sex expression, and fertility. |
| | GO:0009725 | Response to hormone | Broad hormone response; useful as a secondary regulatory category. |
| | GO:0009733 | Response to auxin | Auxin response; relevant to organ initiation, patterning, and reproductive development. |
| | GO:0009737 | Response to abscisic acid | ABA response; relevant to stress-linked fertility, senescence, and developmental regulation. |
| | GO:0009739 | Response to gibberellin | Gibberellin response; often connected to floral development and fertility regulation. |
| | GO:0009740 | Gibberellic acid mediated signaling pathway | GA signaling; often connected to floral development and fertility regulation. |
| | GO:0009735 | Response to cytokinin | Cytokinin response; relevant to meristem activity, organ development, and reproductive growth. |
| | GO:0009723 | Response to ethylene | Ethylene response; relevant to sex expression, senescence, and abscission in some plants. |
| | GO:0009873 | Ethylene-activated signaling pathway | Ethylene signaling; relevant to floral sex expression, senescence, and abscission in some systems. |
| | GO:0009753 | Response to jasmonic acid | Jasmonate response; relevant to anther development, pollen fertility, and defense-fertility tradeoffs. |
| | GO:0009867 | Jasmonic acid mediated signaling pathway | Jasmonate signaling; relevant to anther development, pollen fertility, and stress responses. |
| | GO:0009751 | Response to salicylic acid | Salicylic acid response; broad defense/stress hormone term that may interact with fertility. |
| | GO:0080167 | Response to karrikin | Karrikin response; secondary signaling term, potentially relevant to developmental regulation. |
| Transcriptional/epigenetic | GO:0006355 | Regulation of DNA-templated transcription | Broad transcriptional regulation; useful for transcription factors and sex-linked regulators. |
| | GO:0006357 | Regulation of transcription by RNA polymerase II | Broad transcriptional regulation by Pol II; useful for candidate regulatory genes. |

| Category | GO ID | GO Term | Relevance |
|---|---|---|---|
| | GO:0045892 | Negative regulation of DNA-templated transcription | Transcriptional repression; relevant to regulatory suppression of developmental programs. |
| | GO:0045893 | Positive regulation of DNA-templated transcription | Transcriptional activation; relevant to regulatory activation of developmental programs. |
| | GO:0009887 | Animal organ morphogenesis — exclude for plants unless annotation weirdness appears | Caution/exclude for plant analyses unless it appears from annotation artifacts that need flagging. |
| | GO:0006325 | Chromatin organization | Chromatin organization; relevant to broad epigenetic/regulatory mechanisms. |
| | GO:0016568 | Chromatin modification | Chromatin modification; relevant to epigenetic regulation of reproductive development. |
| | GO:0006342 | Chromatin silencing | Chromatin-based gene silencing; relevant to sex-linked or epigenetic regulation. |
| | GO:0031047 | Gene silencing by RNA | RNA-mediated gene silencing; relevant to small-RNA regulatory mechanisms. |
| | GO:0016441 | Posttranscriptional gene silencing | Posttranscriptional gene silencing; relevant to regulatory control of gene expression. |
| | GO:0035194 | Posttranscriptional gene silencing by RNA | RNA-mediated posttranscriptional silencing; relevant to regulatory control of gene expression. |
| | GO:0080188 | Gene silencing by siRNA-directed DNA methylation | Small-RNA-directed DNA methylation; relevant to epigenetic regulation and genome silencing. |
| | GO:0040029 | Regulation of gene expression, epigenetic | Epigenetic regulation of gene expression; relevant to stable regulatory changes in sex expression. |

Supplementary Table S7. Chloroplast genome assembly statistics for the *Amaranthus* accessions assembled with GetOrganelle v1.7.7.1 and annotated with PGA v2 via PlastidHub v1.0, plus downloaded outgroup plastomes.

| Tip name | Source | Accession | Size (bp) | GC (%) | CDS | tRNA | rRNA |
|---|---|---|---|---|---|---|---|
| *A. acanthochiton* M SRA2 | SRA | SRR19158647 | 150,653 | 36.59 | 81 | 29 | 4 |
| *A. acanthochiton* F SRA1 | SRA | SRR19158648 | 150,653 | 36.59 | 81 | 29 | 4 |
| *A. arenicola* SRA1 | SRA | SRR19158645 | 150,655 | 36.61 | 81 | 29 | 4 |
| *A. australis* M SRA1 | SRA | SRR19158644 | 150,011 | 36.62 | 81 | 29 | 4 |
| *A. cannabinus* M USDA 2021 | This study | SRR39067037 | 150,677 | 36.56 | 81 | 29 | 4 |
| *A. cannabinus* M SRA2 | SRA | SRR19158642 | 150,677 | 36.56 | 81 | 29 | 4 |
| *A. cannabinus* F SRA1 | SRA | SRR19158643 | 150,677 | 36.56 | 81 | 29 | 4 |
| *A. caudatus* USDA 2021 | This study | SRR39067040 | 150,757 | 36.57 | 81 | 29 | 4 |
| *A. cruentus* USDA 2021 | This study | SRR39067039 | 150,757 | 36.57 | 81 | 29 | 4 |
| *A. fimbriatus* USDA 2021 | This study | SRR39067046 | 150,810 | 36.51 | 81 | 29 | 4 |
| *A. floridanus* M SRA1 | SRA | SRR19158641 | 150,670 | 36.60 | 81 | 29 | 4 |
| *A. greggii* USDA AGF1 | This study | SRR39067044 | 150,735 | 36.58 | 81 | 30 | 4 |
| *A. greggii* USDA AGM11 | This study | SRR39067043 | 150,735 | 36.58 | 80 | 30 | 4 |
| *A. hybridus* SRA1 | SRA | SRR12075659 | 150,798 | 36.56 | 81 | 30 | 4 |
| *A. hybridus* SRA2 | SRA | SRR14055740 | 150,759 | 36.56 | 81 | 29 | 4 |
| *A. hypochondriacus* SRA1 | SRA | SRR2130053 | 150,759 | 36.56 | 81 | 30 | 4 |
| *A. hypochondriacus* SRA2 | SRA | SRR2130055 | 150,759 | 36.57 | 81 | 29 | 4 |
| *A. palmeri* F USDA P04 | This study | SRR39067036 | 150,717 | 36.60 | 81 | 30 | 4 |
| *A. palmeri* M USDA P12 | This study | SRR39067035 | 150,717 | 36.60 | 81 | 30 | 4 |
| *A. pumilus* DS002 | This study | SRR39067042 | 150,445 | 36.59 | 81 | 29 | 4 |
| *A. pumilus* EP001 | This study | SRR39067041 | 150,425 | 36.60 | 81 | 29 | 4 |
| *A. retroflexus* USDA 2021 | This study | SRR39067045 | 150,709 | 36.59 | 81 | 29 | 4 |
| *A. spinosus* USDA 2021 | This study | SRR39067038 | 150,524 | 36.61 | 81 | 29 | 4 |
| *A. spinosus* SRA1 | SRA | SRR7121582 | 150,524 | 36.61 | 81 | 30 | 4 |
| *A. tricolor* SRA1 | SRA | SRR17777275 | 150,718 | 36.56 | 81 | 29 | 4 |
| *A. tricolor* SRA2 | SRA | SRR21968681 | 150,717 | 36.56 | 81 | 30 | 4 |

| Tip name | Source | Accession | Size (bp) | GC (%) | CDS | tRNA | rRNA |
|---|---|---|---|---|---|---|---|
| *A. tuberculatus* F USDA T02 | This study | SRR39067034 | 150,673 | 36.60 | 81 | 30 | 4 |
| *A. tuberculatus* M USDA SD008 | This study | SRR39067033 | 150,673 | 36.60 | 81 | 29 | 4 |
| *A. viridis* SRA1 | SRA | SRR7121710 | 150,310 | 36.64 | 81 | 29 | 4 |
| *A. watsonii* F SRA1 | SRA | SRR19158638 | 150,706 | 36.61 | 81 | 29 | 4 |
| *A. watsonii* M SRA2 | SRA | SRR19158646 | 150,706 | 36.61 | 81 | 30 | 4 |
| *Alt. philoxeroides* NCBI ref | NCBI RefSeq | NC_042798.1 | 152,255 | 36.40 | 77 | 19 | 4 |
| *D. amaranthoides* NCBI ref | NCBI RefSeq | NC_041267.1 | 155,108 | 36.78 | 78 | 29 | 4 |

Supplementary Table S8. Mitochondrial reference genomes used to build the per-locus alignment panels for the mitochondrial dataset. Five *Amaranthus* mitochondrial genomes contributed full-length locus copies, and *Alternanthera philoxeroides* served as the direct outgroup reference.

| **Reference** | **NCBI accession** | **Annotated mt features** |
|---|---:|---:|
| *A. hypochondriacus* | PX126027.1 | 34 |
| *A. palmeri* | PX126026.1 | 32 |
| *A. retroflexus* | PX126025.1 | 33 |
| *A. tricolor* | NC_086685.1 | 14 |
| *A. tuberculatus* | CM136843.1 | 44 |
| *Alt. philoxeroides* | MN166292.1 | 26 |

Supplementary Table S9. Mitochondrial locus recovery for samples retained in the mitochondrial supermatrix. *Alternanthera philoxeroides* and *Celosia argentea* were mapped against the outgroup mitochondrial reference panel directly.

| Tip name | Source | Accession | Loci recovered (of 34) | Mean depth |
|---|---|---|---|---|
| *A. acanthochiton* F SRA1 | SRA | SRR19158648 | 34 / 34 | 616.14 |
| *A. acanthochiton* M SRA2 | SRA | SRR19158647 | 34 / 34 | 588.05 |
| *A. arenicola* SRA1 | SRA | SRR19158645 | 34 / 34 | 751.03 |
| *A. australis* M SRA1 | SRA | SRR19158644 | 34 / 34 | 669.88 |
| *A. cannabinus* F SRA1 | SRA | SRR19158643 | 34 / 34 | 887.67 |
| *A. cannabinus* M USDA 2021 | This study | SRR39067037 | 34 / 34 | 507.49 |
| *A. cannabinus* M SRA2 | SRA | SRR19158642 | 34 / 34 | 456.77 |
| *A. caudatus* USDA 2021 | This study | SRR39067040 | 34 / 34 | 480.47 |
| *A. cruentus* USDA 2021 | This study | SRR39067039 | 34 / 34 | 318.48 |
| *A. fimbriatus* USDA 2021 | This study | SRR39067046 | 34 / 34 | 757.21 |
| *A. floridanus* M SRA1 | SRA | SRR19158641 | 34 / 34 | 849.64 |
| *A. greggii* USDA AGF1 | This study | SRR39067044 | 34 / 34 | 705.39 |
| *A. greggii* USDA AGM11 | This study | SRR39067043 | 34 / 34 | 522.72 |
| *A. hybridus* SRA1 | SRA | SRR12075659 | 34 / 34 | 987.88 |
| *A. hybridus* SRA2 | SRA | SRR14055740 | 34 / 34 | 639.12 |
| *A. hypochondriacus* SRA1 | SRA | SRR2130053 | 34 / 34 | 706.53 |
| *A. hypochondriacus* SRA2 | SRA | SRR2130055 | 34 / 34 | 626.41 |
| *A. palmeri* F USDA P04 | This study | SRR39067036 | 34 / 34 | 528.76 |
| *A. palmeri* M USDA P12 | This study | SRR39067035 | 34 / 34 | 581.72 |
| *A. pumilus* DS002 | This study | SRR39067042 | 34 / 34 | 1089.58 |
| *A. pumilus* EP001 | This study | SRR39067041 | 34 / 34 | 1058.08 |
| *A. retroflexus* USDA 2021 | This study | SRR39067045 | 34 / 34 | 267.59 |
| *A. spinosus* USDA 2021 | This study | SRR39067038 | 34 / 34 | 546.23 |
| *A. spinosus* SRA1 | SRA | SRR7121582 | 34 / 34 | 536.76 |
| *A. tricolor* SRA2 | SRA | SRR21968681 | 33 / 34 | 302.26 |

| Tip name | Source | Accession | Loci recovered (of 34) | Mean depth |
|---|---|---:|---:|---:|
| *A. tuberculatus* F USDA T02 | This study | SRR39067034 | 34 / 34 | 300.21 |
| *A. tuberculatus* M USDA SD008 | This study | SRR39067033 | 34 / 34 | 372.64 |
| *A. viridis* SRA1 | SRA | SRR7121710 | 34 / 34 | 357.74 |
| *A. watsonii* F SRA1 | SRA | SRR19158638 | 34 / 34 | 604.01 |
| *A. watsonii* M SRA2 | SRA | SRR19158646 | 34 / 34 | 646.12 |
| *Alt. philoxeroides* SRA1 | SRA | ERR6752173 | 31 / 34 | — |
| *C. argentea* SRA1 | SRA | SRR15412865 | 28 / 34 | — |

Supplementary Table S10. Penalized-likelihood ultrametric model comparison for the rooted ASTRAL nuclear species tree pruned to the *Amaranthus* ingroup. Note: models were statistically indistinguishable in fit, and the strict clock was retained as the most parsimonious model.

| **Method** | **Parameter** | **Log-likelihood** | **PHIIC** |
|---|---|---|---|
| Clock (strict) | — | -0.879 | 73.758 |
| Discrete | k = 1 | -0.879 | 73.758 |
| Discrete | k = 2 | -0.879 | 77.758 |
| Discrete | k = 3 | -0.879 | 81.758 |
| Discrete | k = 4 | -0.879 | 85.758 |
| Discrete | k = 5 | -0.879 | 89.758 |
| Correlated | λ = 0.01 | -0.873 | 215.746 |
| Correlated | λ = 0.1 | -0.875 | 215.748 |
| Correlated | λ = 0.5 | -0.876 | 215.751 |
| Correlated | λ = 1.0 | -0.876 | 215.752 |
| Correlated | λ = 2.0 | -0.877 | 215.753 |
| Correlated | λ = 5.0 | -0.877 | 215.754 |
| Correlated | λ = 10.0 | -0.877 | 215.754 |
| Relaxed | λ = 1.0 | -0.884 | 215.768 |
